# Molecular basis of sarbecovirus evolution and receptor tropism in natural hosts, potential intermediate hosts, and humans

**DOI:** 10.1101/2025.03.22.644775

**Authors:** Yusuke Kosugi, Kanata Matsumoto, Spyros Lytras, Arnon Plianchaisuk, Jarel Elgin Tolentino, Shigeru Fujita, Maximilian Stanley Yo, Chen Luo, Yoonjin Kim, Wataru Shihoya, Jumpei Ito, Osamu Nureki, Kei Sato

## Abstract

The spike protein of many sarbecoviruses binds to the angiotensin-converting enzyme 2 (ACE2) receptor and facilitates viral entry. The diversification of the sarbecovirus *spike* gene and the mammalian *ACE2* gene suggests that sarbecoviruses and their hosts have co-evolved, and the genetic diversity in these genes affects the host tropism of sarbecoviruses. Better comprehending the evolutionary potential of sarbecoviruses can lead to preparedness for the next pandemic. However, the host tropism of sarbecoviruses is not fully understood. Here, we performed round-robin pseudovirus infection assays using 53 sarbecoviruses and ACE2s from 17 mammals to elucidate the ACE2 tropism of sarbecoviruses in natural hosts, potential intermediate hosts and humans. We determined the factors responsible for the ACE2 tropism of sarbecoviruses through structural, phylogenetic analyses, and infection experiments, revealing which substitutions can expand the host range of sarbecoviruses. These results highlight the mechanisms modulating host tropism throughout sarbecovirus evolution.

## Introduction

Many pathogenic and contagious viruses that have emerged in documented human history, such as Ebola virus, Nipah virus, and human immunodeficiency virus, originated from the spillover of viruses carried by wildlife to humans. Current scientific evidence supports that SARS-CoV-2, the causative agent of the COVID-19 pandemic, is another example of a human pathogenic virus emerging through zoonotic transmission^1–3^.

In recent years, many coronaviruses phylogenetically close to SARS-CoV or SARS-CoV-2 (members of the subgenus *Sarbecovirus*) have been reported, most in East and Southeast Asia^4–8^. Reservoir hosts of sarbecoviruses are *Rhinolophus* bats found in East and Southeast Asia, and these viruses have also been found in the same bat genus in Africa and Europe^9–11^. Other than *Rhinolophus* bats, a handful of sarbecoviruses have been found in wild or captive pangolins, although these likely represent independent spillovers from bats^12–14^.

The spike (S) glycoprotein of sarbecoviruses binds to the host receptor and facilitates viral entry into cells. Many sarbecoviruses, including SARS-CoV and SARS-CoV-2, use the angiotensin-converting enzyme 2 (ACE2) as their entry receptor, while the receptor of some bat sarbecoviruses is still unknown^15,16^. The receptor-binding domain (RBD) of sarbecovirus S proteins is highly diversified into distinct clades corresponding to unique functional and structural characteristics^17,18^. Similarly, mammalian *ACE2* genes are highly diversified^19–21^. This genetic diversity in both virus and host results in a complex host tropism that determines which mammalian hosts each sarbecovirus can potentially infect.

It is suggested that sarbecoviruses undergo repeated inter-species transmissions between different *Rhinolophus* species in Southeast Asia, and that the multiple transmission events have facilitated the evolution of sarbecoviruses^4,7,12,13,22^. Previous studies including ours have focused on the tropism of sarbecovirus S proteins to various mammalian ACE2 proteins and addressed the potential host range of sarbecoviruses^17,18,23–25^. These previous studies focused on a small subset of sarbecoviruses related to human sarbecoviruses (i.e., SARS-CoV or SARS-CoV-2), such as WIV1, RaTG13 and GX-P2V, and showed how they can utilise the ACE2 proteins of some bats and other mammals for cellular entry. On the other hand, there is a number of sarbecovirus members that do not seem to use any of the mammalian ACE2 proteins tested^23,24^. To date, more than 200 sarbecoviruses have been identified. However, the number of sarbecovirus S and mammalian ACE2s used in previous studies is limited, and the full range of host tropism across the sarbecoviruses is not fully understood.

The fact that two sarbecoviruses, SARS-CoV and SARS-CoV-2, have emerged and caused outbreaks in the past two decades suggests that similar spillover events potentially leading to a pandemic may occur in the future. Understanding the evolutionary potential of sarbecoviruses circulating in *Rhinolophus* bats, putative intermediate hosts, and humans can aid our preparedness for the next pandemic. In this study, we elucidated the ACE2 tropism of a wide range of sarbecoviruses by performing a round-robin pseudovirus infection assays using 53 sarbecovirus S proteins and 17 ACE2 proteins of natural hosts (*Rhinolophus* bats and pangolins), potential intermediate hosts (civets and raccoon dogs), and humans. We then integrated protein structural analyses, phylogenetic approaches, and infection experiments to determine the factors responsible for the ACE2 tropism of sarbecoviruses.

## Results

### The spectrum of sarbecovirus ACE2 tropism

We set out to understand the phylogenetic relationship between the known sarbecoviruses by focusing on their RBD sequences, which should be the key determinant of their receptor tropism. Consistent with previous reports^17,18^, we categorized sarbecoviruses based on their RBDs into four clades: Clade 1a (SARS-CoV-related sarbecoviruses), Clade 1b (SARS-CoV-2-related sarbecoviruses), Clade 2 (non-ACE2-usage sarbecoviruses), and Clade 3 (European/African sarbecoviruses) (**Figures 1A and S1**). Among the 254 sarbecoviruses listed in **Table S1**, we selected 53 sarbecoviruses from most of the subclades in each clade for the experiment (summarized in **Figure S1**).

**Figure 1.**
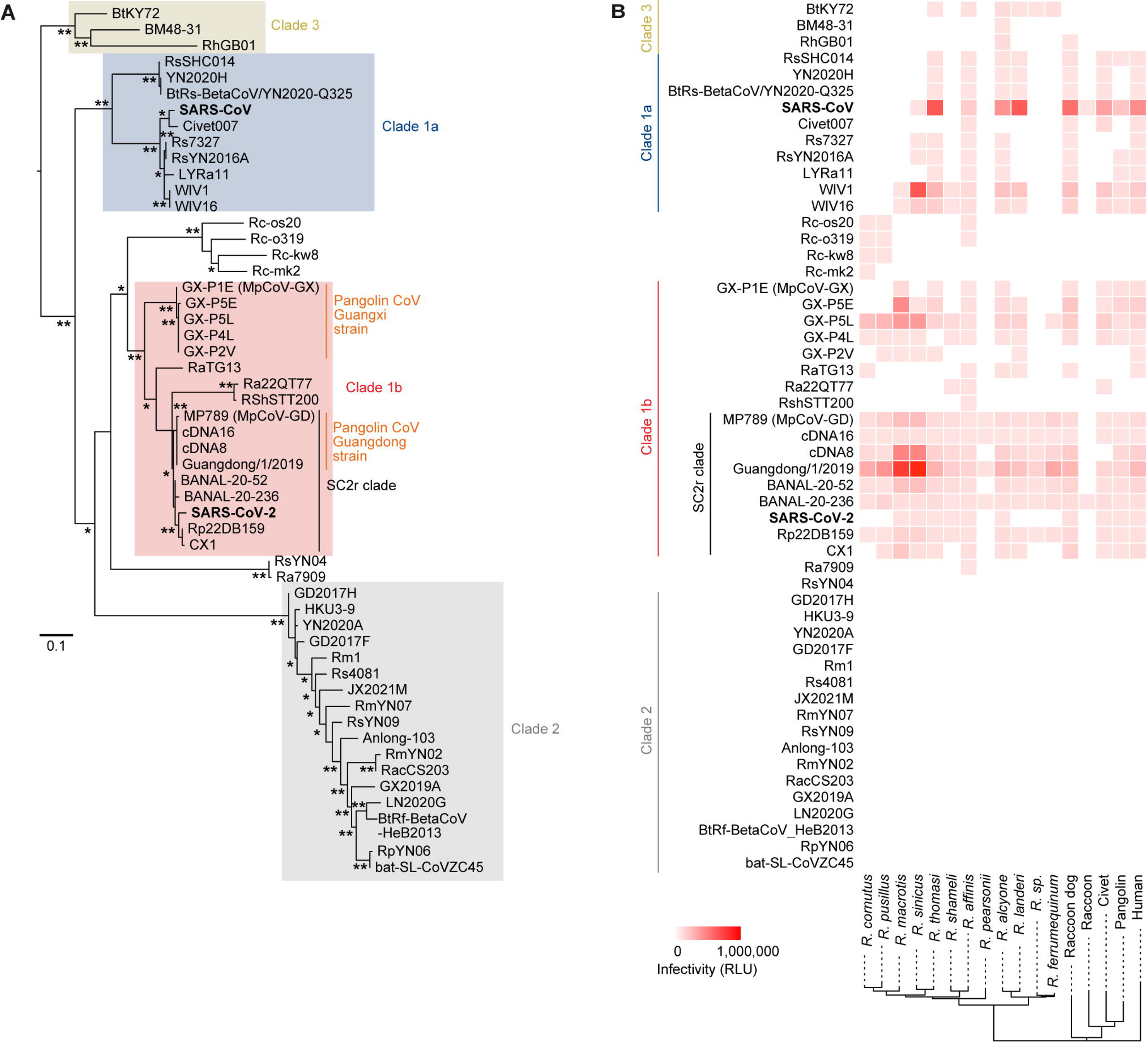
Round-robin analysis of the ACE2 tropism of 53 sarbecoviruses. (A) Maximum likelihood tree of the 53 sarbecoviruses used for pseudovirus assays based on their S RBD amino acid sequences. *, bootstrap value >80; **, bootstrap value >90. Scale bar indicates genetic distance (amino acid substitutions per site). The names of sarbecovirus clades (Clade 1a, Clade 1b, Clade 2 and Clade 3) highlighted in the tree correspond to a previous work^18^. SC2r clade, SARS-CoV-2-related clade. (B) Round-robin pseudovirus assay. HIV-1-based reporter viruses pseudotyped with the S proteins of 53 sarbecoviruses were prepared. The pseudoviruses were inoculated into a series of HOS-TMPRSS2 cells stably expressing *Rhinolophus* bat ACE2. The heatmap shows representative mean values of pseudovirus infectivity (quadruplicates of independently bound or infected cells) from three independent experiments. Related to **Figures S1 and S2** and **Tables S1 and S2**.

To investigate the host ACE2 tropism of sarbecoviruses, we prepared a series of human HOS-TMPRSS2 cell lines stably expressing ACE2 proteins from one of twelve *Rhinolophus* bat species: *R. affinis, R. cornutus, R. ferrumequinum, R. macrotis, R. pearsonii, R. pusillus, R. shameli, R. alcyone, R. landeri, R. thomasi, R. sinicus*, and an unidentified *Rhinolophus* species from Uganda (hereafter referred to as “*R. sp*.”) ^26^. The cells stably expressing ACE2 proteins of human, raccoon dog, raccoon, civet, and pangolin were also prepared. We then prepared lentivirus-based pseudoviruses with the S proteins of 53 sarbecoviruses and inoculated them into a series of ACE2-expressing target cells. As shown in **Figures 1B and S2**, not all sarbecoviruses can use human ACE2 as a receptor. Consistent with a previous study^27^, almost none of the sarbecoviruses could use raccoon ACE2 (**Figures 1B and S2**). Moreover, Clade 2 sarbecoviruses did not show any infectivity in the cells expressing any ACE2 proteins tested (**Figures 1B and S2**), consistent with previous studies^17,18,23,24^.

Among Clade 3 viruses, although BtKY72 showed infectivity in the cells expressing ACE2 of several *Rhinolophus* species, including *R. thomasi*, *R. affinis*, *R. alcyone*, *R. landeri*, *R. sp*., and *R. ferrrumequinum*, BM48-31 and RhGB01 only infected the cells expressing *R. alcyone* ACE2 (**Figures 1B and S2**). All sarbecoviruses in Clade 1a showed infectivity in the cells expressing ACE2 proteins from multiple *Rhinolophus* species but also those from raccoon dog, civet, pangolin, and human. All sarbecoviruses in Clade 1b, except for RshSTT200 and Ra22QT77, showed infectivity in cells expressing human ACE2 (**Figures 1B and S2**).

Four sarbecoviruses detected in *R. cornutus* in Japan^28,29^, Rc-o319, Rc-os20, Rc-mk2, and Rc-kw8, are phylogenetically close to Clade 1b viruses (**Figures 1B and S2**). However, these four pseudoviruses did not show infectivity in human ACE2-expressing cells, but only infected the cells expressing the ACE2 of a few *Rhinolophus* species, including their host, *R. cornutus* (**Figures 1B and S2**). Altogether, these results suggest that the ACE2 tropism of sarbecoviruses can be mainly defined by the RBD clade, although some viruses have a unique tropism profile compared to the other viruses in the same clade.

### Residues enabling BtKY72 S to use human ACE2

BtKY72 S exhibited infectivity in *R. ferrumequinum* ACE2 but did not in human ACE2 (**Figure 1B**). In contrast, SARS-CoV-2 S exhibited infectivity in human ACE2 but did not in *R. ferrumequinum* ACE2 (**Figure 1B**). Since previous studies have suggested the importance of the residues positioned at 493 and 498 of BtKY72 S protein on the usage of human ACE2^23^, we focused on these two residues on BtKY72 and SARS-CoV-2: BtKY72 S harbors K493 and T498, whereas SARS-CoV-2 S harbors Q493 and Q498. To address whether the interaction between the amino acid residues 493 and 498 in S and the corresponding residues in ACE2 determine the difference in ACE2 tropism between BtKY72 and SARS-CoV-2, we prepared the S derivatives of BtKY72 harboring the respective substitutions at residues 493 and 498. We then performed infectivity assays using the cells expressing ACE2s from eight *Rhinolophus* bats, pangolin, and human. As shown in **Figure 2A**, the BtKY72 S possessing K493Q and T498Q exhibited infectivity to cells expressing human ACE2. On the other hand, these substitutions did not confer infectivity to the cells expressing ACE2 of *R. cornutus, R. shameli, R. pearsonii, R. pusillus, R. macrotis*, and *R. sinicus*, and only the BtKY72 S possessing T498Q showed infectivity in *R. affinis* ACE2-expressing cells (**Figure 2A**). Interestingly, although the BtKY72 S with K493Q/T498Q substitutions conferred infectivity in human ACE2-expressing cells, this derivative lost its ability to infect *R. ferrumequinum* ACE2-expressing cells (**Figure 2A**). These results suggest that the usage of human ACE2 and *R. ferrumequinum* ACE2 is mutually exclusive for the BtKY72 S and is primarily determined by the amino acid residues at positions 493 and 498.

**Figure 2.**
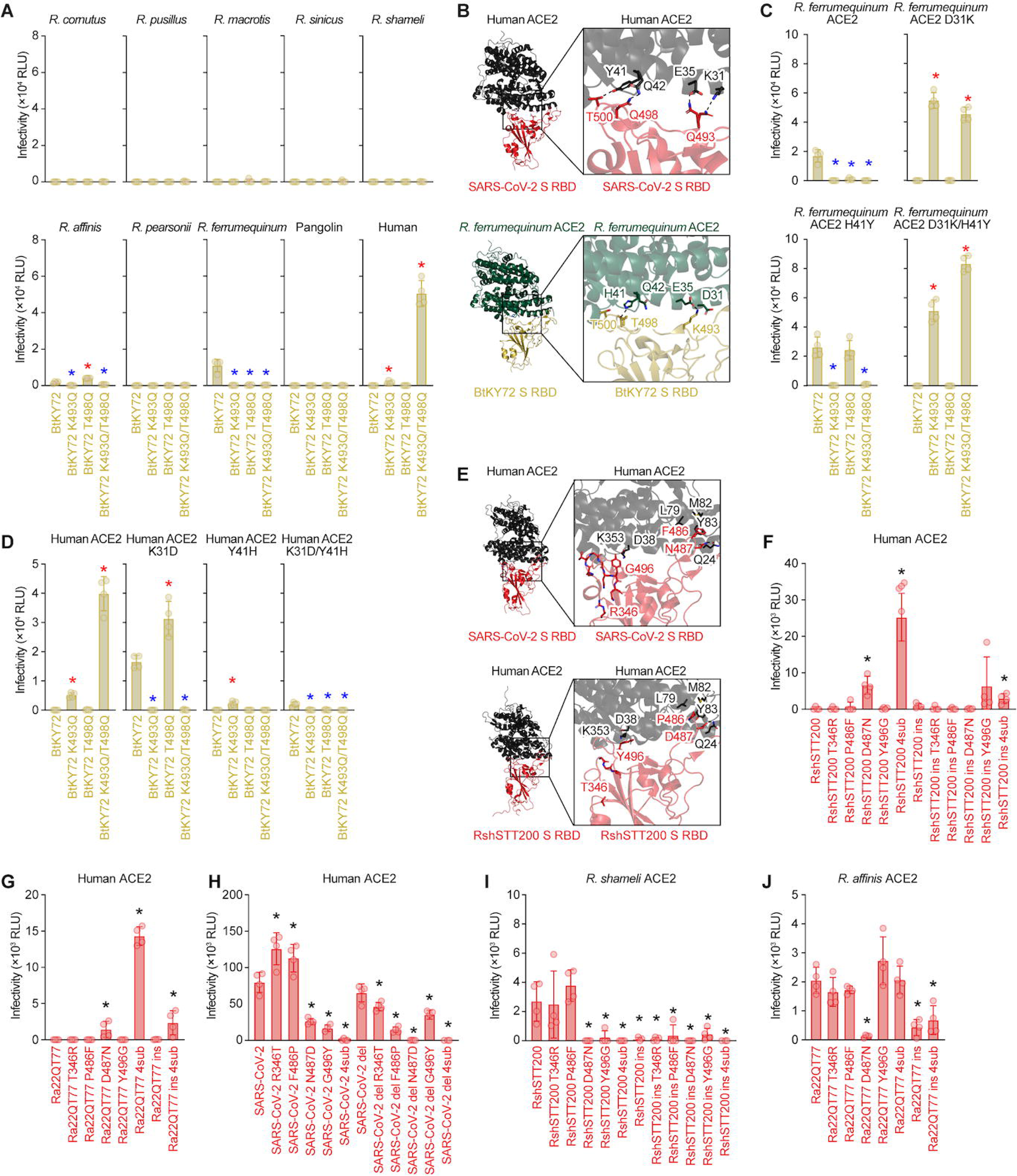
Identification of amino acid residues in S protein determining human ACE2 tropism (A) Pseudovirus assay. HIV-1-based reporter viruses pseudotyped with the S proteins of BtKY72 or their derivatives were prepared. The pseudoviruses were inoculated into a series of HOS-TMPRSS2 cells stably expressing *Rhinolophus* bat ACE2. (B) Structural insights into the binding of S RBD and ACE2 proteins. (top) The structural model of complex of SARS-CoV-2 S RBD (red) and the human ACE2 (black) (PDB:6M0J)^30^. (bottom) The structural model of complex of BtKY72 S RBD (yellow) and *R. ferrumequinum* ACE2 modeled by AlphaFold3 (green). The model was reconstructed by using co-structure of BtKY72 S RBD and *R. landeri* ACE2 (PDB:8K4U)^31^ as template, and models predicted by AlphaFold3. Residues 493 and 498 of S RBD and residues 31, 35, 41 and 42 of ACE2s are indicated as stick model. Dashed lines indicate hydrogen bonds. The residue numbering is based on SARS-CoV-2. (**C-D**) Pseudovirus assay. HIV-1-based reporter viruses pseudotyped with the S proteins of BtKY72 or their derivatives were prepared. The pseudoviruses were inoculated into a series of HEK293 cells transiently expressing *Rhinolophus* bat ACE2. (E) (top) The structural model of complex of SARS-CoV-2 S RBD (red) and the human ACE2 (black) (PDB:6M0J)^30^. (bottom) The structural model of complex of RshSTT200 S RBD (red) and the human ACE2 (black) (PDB:7XBH)^34^. Residues of S RBD with substitutions or deletions focused on in this study and residues of ACE2s forming interaction with these are indicated as stick models. Dashed lines indicate hydrogen bonds. The residue numbering is based on SARS-CoV-2. (**F-H**) Pseudovirus assay. HIV-1-based reporter viruses pseudotyped with the S proteins of (**F**) RshSTT200, (**G**) Ra22QT77, (**H**) SARS-CoV-2 or their derivatives were prepared. The pseudoviruses were inoculated into a series of HOS-TMPRSS2 cells stably expressing human ACE2. (I) Pseudovirus assay. HIV-1-based reporter viruses pseudotyped with the S proteins of RshSTT200 or their derivatives were prepared. The pseudoviruses were inoculated into a series of HOS-TMPRSS2 cells stably expressing *R. shameli* ACE2. (J) Pseudovirus assay. HIV-1-based reporter viruses pseudotyped with the S proteins of Ra22QT77 or their derivatives were prepared. The pseudoviruses were inoculated into a series of HOS-TMPRSS2 cells stably expressing *R. affinis* ACE2. In (**A**), (**C**)-(**D**), and (**F**)-(**J**), the infectivity (relative light unit) in each target cell is shown. Data are expressed as the mean with SD. Assays were performed in triplicate. The number in the panel indicates the fold change versus parental S. Statistically significant differences (*, *P* < 0.05) between wildtype and the derivatives were determined by two-sided Student’s t test. Red and blue asterisks show the significantly higher and lower infectivity than wildtype, respectively. Related to **Figure S3** and **Table S3**.

To determine the amino acid residues in ACE2 that are responsible for the mutually exclusive usage of human ACE2 and *R. ferrumequinum* ACE2 by the BtKY72 S, the amino acid residues in ACE2 that potentially interact with the residues 493 and 498 in S were assessed using the co-structure of SARS-CoV-2 S and human ACE2 (PDB:6M0J)^30^. This co-structure showed that the residue 493 in S interacts with the residues 31 and 35 in human ACE2, while the residue 498 in S interacts with the residues 41 and 42 in human ACE2 (**Figure 2B**). Similarly, a previous paper showed that the residue 493 in BtKY72 S interacts with residues 31 and 35 in *R. ferrumequinum* ACE2, while the residue 498 in S interacts with the residue 41 in *R. ferrumequinum* ACE2^31^. When comparing the amino acid sequences of human ACE2 and *R. ferrumequinum* ACE2, the residues at the corresponding positions at 35 (E) and 42 (Q) were identical (**Figure 2B**). However, human ACE2 has K31 and Y41, while *R. ferrumequinum* ACE2 has D31 and H41 (**Figure 2B**). To further address whether the difference in the residues positioned at 31 and/or 41 in ACE2 is responsible for the ACE2 usage of BtKY72 S, we prepared the plasmids expressing *R. ferrumequinum* ACE2 and human ACE2 that swapped the residues positioned at 31 and 41. As shown in **Figure 2C**, the D31K substitution reverted the use of *R. ferrumequinum* ACE2 by the BtKY72 S. Consistent with previous studies^32,33^, the BtKY72 S with T498Q showed infectivity in the cells expressing *R. ferrumequinum* ACE2 with H41Y (**Figure 2C**).

In the case of human ACE2, the K31D substitution conferred susceptibility to BtKY72 S (**Figure 2D**). Although the BtKY72 S with T498Q showed infectivity in the cells expressing the human ACE2 with K31D, this infectivity was canceled when the Y41H substitution was introduced (**Figure 2D**). These results suggest that the mutually exclusive use of *R. ferrumequinum* ACE2 and human ACE2 by the BtKY72 S is determined by two intermolecular interactions: one between the residue 493 in S and the residue 31 in ACE2; and another between the residue 498 in S and the residue 41 in ACE2.

### Residues preventing certain Clade 1b sarbecoviruses from using human ACE2

RshSTT200 and Ra22QT77 are the only Clade 1b viruses tested in this study that are unable to use human ACE2 for infection (**Figure 1B**). Compared with the SARS-CoV-2 S, the S proteins of RshSTT200 and Ra22QT77 have two deletions in the residues 444-445 and 448-449 (relative to the SARS-CoV-2 S) and four amino acid differences (T346R, P486F, D487N, and Y496G) (**Figure S3A**). A previous study reported that a RshSTT200 S RBD, which filled in these two deletions and contained four SARS-CoV-2 S-derived substitutions, confers the ability to bind to human ACE2^34^. As shown in **Figure 2E**, both the deletions and substitutions are located in the ACE2 binding interface of S. However, the factors (i.e., deletion and/or substitution) that determine the inability of the S proteins of RshSTT200 and Ra22QT77 to use human ACE2 remain unclear. To address this issue, we generated a series of derivatives of RshSTT200 and Ra22QT77 S. In the infectivity assay using the cells expressing human ACE2 and the pseudoviruses with derivatives of RshSTT200 S, we found that a derivative containing four substitutions at positions 346, 486, 487, and 496 (“RshSTT200 4sub” in the figure) shows higher infectivity, while a derivative filling in the two deletions (“RshSTT200 ins” in the figure) does not (**Figure 2F**). These data suggest that the inability to use human ACE2 is determined by the four substitutions but not by the two deletions. Among these four substitutions, a sole substitution, D487N, conferred the ability to use human ACE2 for infection (**Figure 2F**). Similar results were observed in the Ra22QT77 S derivatives (**Figure 2G**): the use of human ACE2 for infection was acquired by the four substitutions (“Ra22QT77 4sub” in the figure), and particularly, the D487N substitution significantly increased the infectivity of Ra22QT77 S pseudovirus. On the other hand, the derivative filling in the two deletions (“Ra22QT77 ins” in the figure) did not show infectivity (**Figure 2G**).

To further address the impact of the two deletions and four substitutions on human ACE2 usage, we generated a series of SARS-CoV-2 S derivatives. Consistent with the results of RshSTT200 (**Figure 2F**) and Ra22QT77 (**Figure 2G**), introducing the two deletions observed in the S proteins of RshSTT200 and Ra22QT77 into the SARS-CoV-2 S (“SARS-CoV-2 del” in the figure) did not affect the infectivity in the cells expressing human ACE2 (**Figure 2H**). Additionally, a derivative with the four substitutions (“SARS-CoV-2 4sub” in the figure) lost the infectivity in human ACE2-expressing cells (**Figure 2H**). Notably, the N487D and G496Y substitutions had the biggest effect on the loss of human ACE2-dependent infectivity in SARS-CoV-2 (**Figure 2H**). These results suggest that acquiring human ACE2 usage for the S proteins of RshSTT200 and Ra22QT77 is determined by the amino acid substitutions positioned at 346, 486, 487, and 496 (487 having the biggest effect), while the two hallmark deletions unique to these viruses are not related to human ACE2 usage.

Since we observed the mutually exclusive use of human ACE2 and *R. ferrumequinum* ACE2 in BtKY72 (**Figures 2A-2D**), we hypothesized that the four substitutions in RshSTT200 and Ra22QT77 S proteins, which confer the ability to utilize human ACE2, may render these S proteins incapable of the use of the ACE2 proteins of their natural hosts (*R. shameli* for RshSTT200, *R. affinis* for Ra22QT77).

In the case of RshSTT200 S (**Figure 2I**), the “4sub” derivative (harboring T346R, P486F, D487N, and Y496G substitutions), which acquired the ability to use human ACE2 for infection (**Figure 2F**), lost the ability to use *R. shameli* ACE2. Particularly, the two substitutions, D487N and Y496G, are responsible for the inability to use *R. shameli* ACE2 (**Figure 2I**). These results suggest that, similar to BtKY72 S, there is a trade-off relationship for RshSTT200 S to use human ACE2 or *R. shameli* ACE2.

In the case of Ra22QT77 S, the D487N derivative lost the infectivity in the cells expressing *R. affinis* ACE2 (**Figure 2J**). However, the “4sub” derivative (harboring T346R, P486F, D487N and Y496G substitutions) of Ra22QT77 S, which acquired the ability to use human ACE2 for infection (**Figure 2G**), showed the infectivity comparable to the parental Ra22QT77 S in *R. affinis* ACE2-expressing cells (**Figure 2J**). These results suggest that the four substitutions positioned at 346, 486, 487 and 496 confer the potential to broaden the receptor usage of Ra22QT77 S to use not only the ACE2 of its natural host, *R. affinis*, but also human ACE2.

Nevertheless, the effect of the D487N substitution in RshSTT200 and Ra22QT77 S on ACE2 preference was mutually exclusive between human and their natural hosts (i.e., *R. shameli* for RshSTT200 and *R. affinis* for Ra22QT77). To assess the effect of the substitution at position 487 on the ACE2 usage of RshSTT200 and Ra22QT77 at the structural level, we examined the ACE2 region structurally proximal to position 487 in S using the co-structure model of SARS-CoV-2 S and human ACE2 (PDB:6M0J)^30^. As shown in **Figure S3B**, we found that the region of ACE2 surrounding residue 487 in S is negatively charged. This observation suggests that this ACE2 region can lead to a repulsive force between itself and the negatively charged D487 of RshSTT200 and Ra22QT77.

### Molecular and structural insights into the host tropism of Rc-o319

Rc-o319 is a sarbecovirus identified in an *R. cornutus* bat in Japan^28^. This virus is unable to use human ACE2 for infection, but can use a few *Rhinolophus* ACE2 proteins, including that of its natural host (**Figures 1B and S2**). To understand the unique interaction between Rc-o319 S and the ACE2 of its natural host, *R. cornutus,* at the structural level, we determined the complex structure of Rc-o319 RBD bound to *R. cornutus* ACE2 by cryo-electron microscopy (**Figures 3A, S3D-S3G and Table S4**). The structure revealed that residues K27, D31, Y41, Q42, E75, and K353 of the *R. cornutus* ACE2 directly interacted with the Rc-o319 S RBD (**Figures 3A and S3H**). The glycosylation of the residue N38 of *R. cornutus* ACE2 was observed, and more interestingly, the N38-linked glycan interacted with the residue N494 of Rc-o319 S RBD (**Figure 3B**).

**Figure 3.**
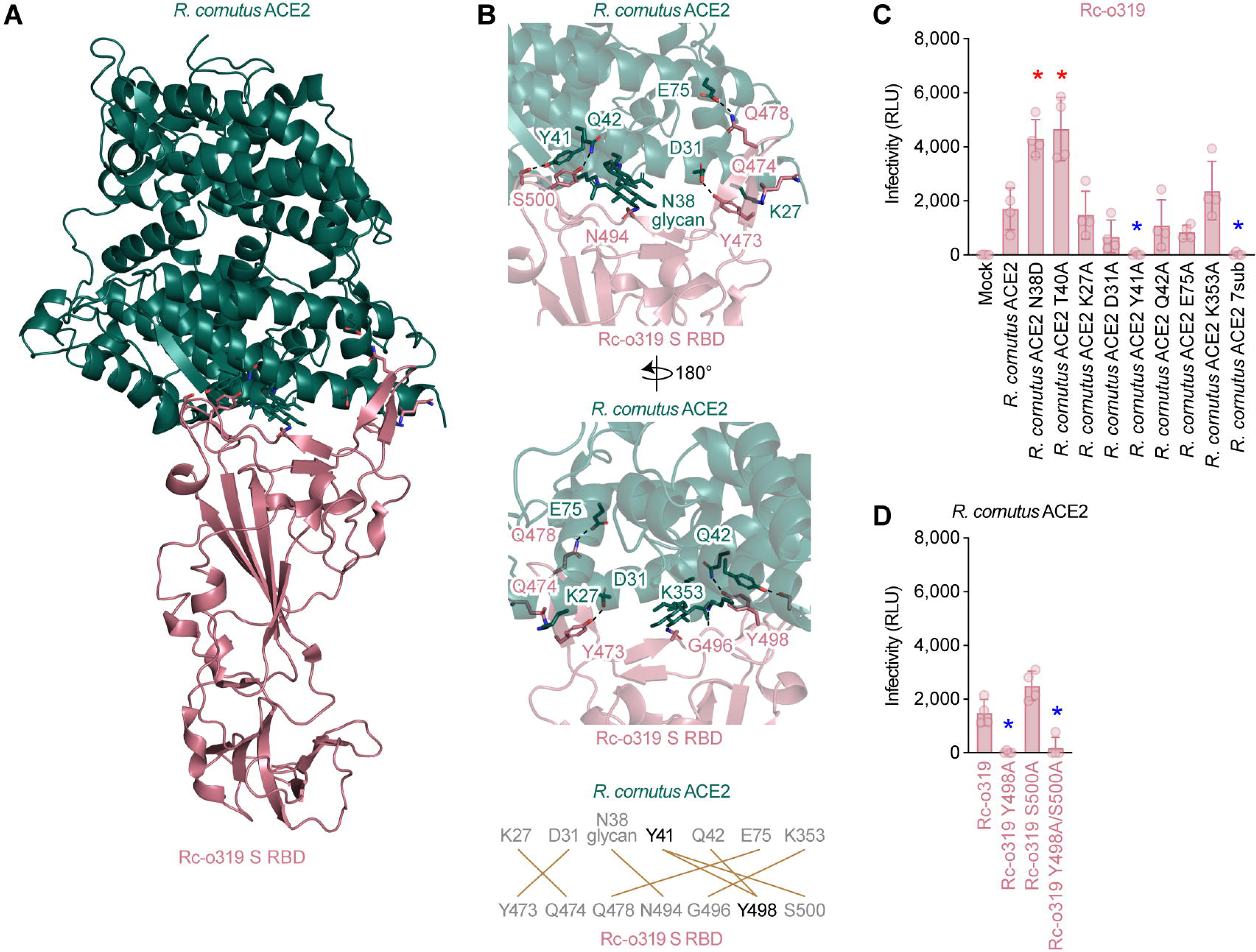
Interaction between Rc-o319 and the *R. cornutus* ACE2 of its natural host (A) (left) The cryo-EM structure of the complex of Rc-o319 S RBD (pink) and *R. cornutus* ACE2 (green). The complex structure is shown as a cartoon, and the residues involved in the hydrogen bond interactions are shown as sticks. (top right) The close-up view of the complex structure. The structure rotated 180° on y axis are also shown on the bottom. Dashed lines represent hydrogen bonds. The residue numbering is based on SARS-CoV-2. (bottom right) The scheme of interaction between Rc-o319 S receptor-binding motif (top) and *R. cornutus* ACE2 (bottom). Salt bridge or hydrogen bond is indicated in brown. Y498 of Rc-o319 S and Y41 of *R. cornutus* ACE2 are indicated in black. RBM, receptor-binding motif. (**B and C**) Pseudovirus assay. HIV-1-based reporter viruses pseudotyped with the S proteins of Rc-o319 or the derivatives were prepared. The pseudoviruses were inoculated into a series of HEK293 cells transiently expressing *R. cornutus* ACE2. the infectivity (relative light unit) in each target cell is shown. Data are expressed as the mean with SD. Assays were performed in triplicate. The number in the panel indicate the fold change versus parental S. Statistically significant differences (*, *P* < 0.05) between wildtype and the derivatives of *R. cornutus* ACE2 (**B**) or Rc-o319 (**C**) were determined by two-sided Student’s t test. Red and blue asterisks show the significantly higher and lower infectivity than wildtype, respectively. Related to **Figure S3** and **Table S4**.

To assess the impact of the *R. cornutus* ACE2 N38-linked glycosylation on binding to the Rc-o319 S RBD, plasmids expressing *R. cornutus* ACE2 with the substitution, N38D or T40A, both of which disrupt the N38-linked glycosylation, were prepared. As shown in **Figure 3C**, the N38D and T40A substitutions significantly increased the infectivity of Rc-o319, suggesting that the N38-linked glycan interferes with the binding of *R. cornutus* ACE2 to Rc-o319 S.

Next, to assess the interaction between *R. cornutus* ACE2 and Rc-o319 S RBD experimentally, we generated plasmids expressing a series of *R. cornutus* ACE2 derivatives in which each residue of interest is substituted with alanine (i.e., K27A, D31A, Y41A, Q42A, E75A, or K353A). We found that the infectivity of Rc-o319 significantly decreased only in the cells expressing *R. cornutus* ACE2 Y41A (**Figure 3C**). Because Y41 of *R. cornutus* ACE2 interacts with Y498 and S500 of Rc-o319 S (**Figure 3B, bottom**), we further generated plasmids expressing three Rc-o319 S derivatives containing Y498A, S500A, or both, and evaluated the importance of these residues in Rc-o319 S for the use of *R. cornutus* ACE2. As shown in **Figure 3D**, Y498A, but not S500A, abolished the infectivity of Rc-o319 in cells expressing *R. cornutus* ACE2. These findings suggest that the interaction between Y498 of Rc-o319 S and Y41 of *R. cornutus* ACE2 is crucial for the binding of Rc-o319 S to *R. cornutus* ACE2.

### Expansion of ACE2 tropism by changing key amino acid residues during sarbecovirus evolution

In **Figure 1B**, some Clade 1b viruses showed a relatively broader range of ACE2 usage. This suggests an expansion of ACE2 tropism during the evolution of Clade 1b sarbecovirus, including SARS-CoV-2. Here we defined this subclade as the “SARS-CoV-2 related (SC2r) clade”. To track the evolution of ACE2 tropism in each sarbecovirus lineage, we calculated the “broadness score” that indicates the ACE2 usage range of each sarbecovirus S protein according to the results of the pseudovirus infection assay shown in **Figure 1B**. We then estimated the broadness scores of sarbecovirus ancestors using a maximum likelihood method ancestral reconstruction method based on their RBD. The ancestral lineage of sarbecoviruses was estimated to have a limited ACE2 usage range (**Figure 4A**).

**Figure 4.**
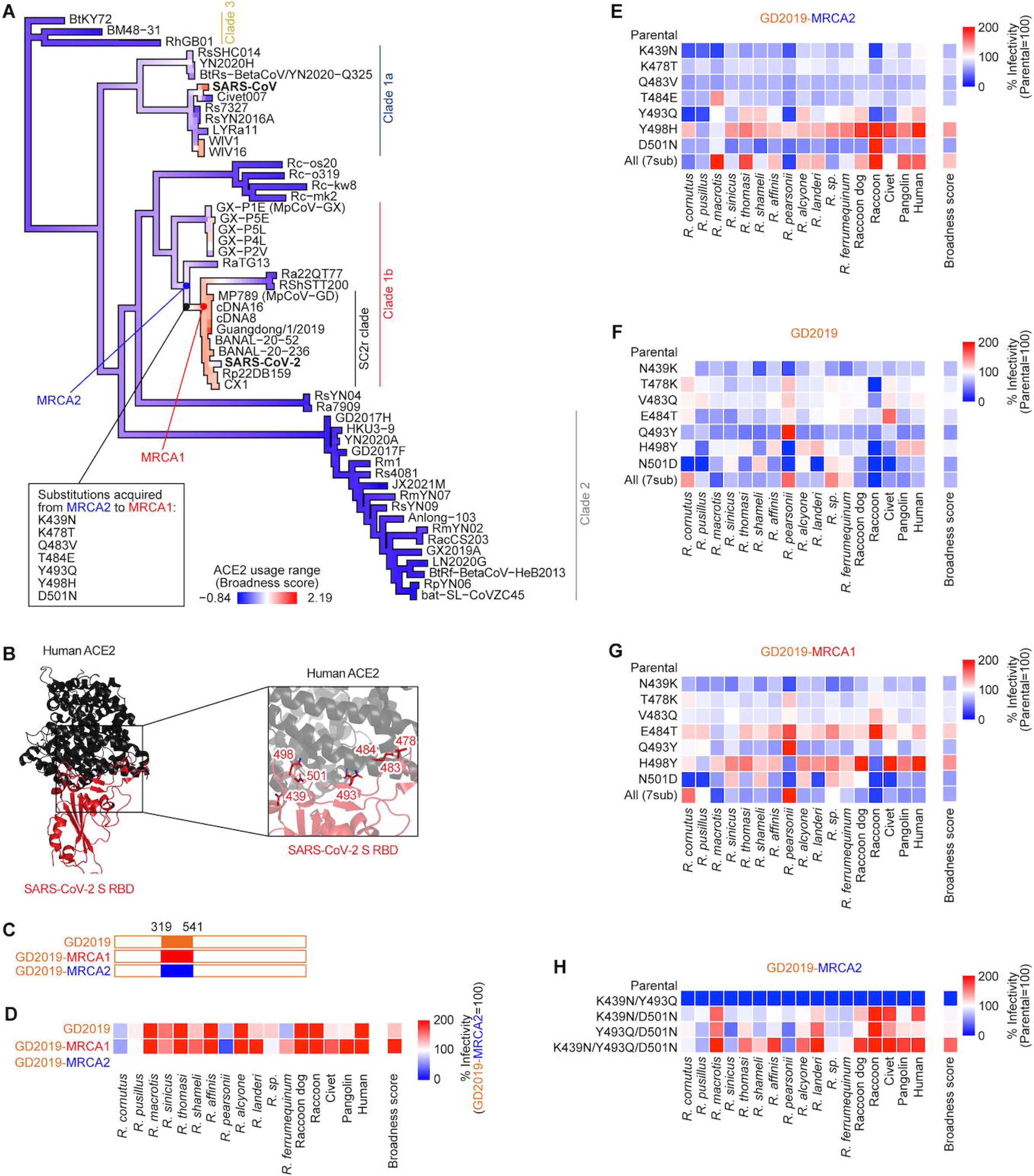
Amino acid substitutions leading to expansion of ACE2 usage range during sarbecovirus evolution (A) The change in ACE2 usage range during the evolution of sarbecoviruses. The color in the phylogenetic tree indicates the ACE2 usage range of each sarbecovirus (defined as the “broadness score”). The broadness score was calculated as the median of the standardized Z score of infectivity in the ACE2-expressing cells. The most recent common ancestor of RaTG13 and a subclade of Clade 1b sarbecoviruses, including SARS-CoV-2 (MRCA2) and that of the subclade (MRCA1) are highlighted. The substitutions that occurred in the branch connecting MRCA2 to MRCA1 are shown in the black box. (B) The structural model of complex of SARS-CoV-2 S RBD (red) and the human ACE2 (black) (PDB:6M0J)^30^. (C) The scheme of Guangdong/1/2019 S (GD2019) and GD2019 replaced the RBD with MRCA1 (GD2019-MRCA1) or MRCA2 (GD2019-MRCA2). (D) Pseudovirus assay. HIV-1-based reporter viruses pseudotyped with the S proteins of with RaTG13, GD2019, GD2019-MRCA2 or GD2019-MRCA1 protein were prepared. (**E-H**) Pseudovirus assay. HIV-1-based reporter viruses pseudotyped with the S proteins of with GD2019-MRCA2 (**E, H**), GD2019 (**F**) or GD2019-MRCA1 (**G**) protein, or the derivatives were prepared. In (**D**)-(**H**), the pseudoviruses were inoculated into a series of HOS-TMPRSS2 cells stably expressing *Rhinolophus* bat ACE2. Heatmap plotted by the mean values of % Infectivity compared to GD2019-MRCA2 (**D**) or parental (**E-H**) (quadruplicates of independently bound or infected cells) which are representative results out of three independent experiments. The broadness score was calculated as the median of % infectivity in the ACE2-expressing cells. Related to **Figure S4** and **Tables S3 and S4**.

When we focus on the ancestors of Clade 1b, the node at the most recent common ancestor (MRCA) of the SC2r clade (“MRCA1” in **Figure 4A**), was estimated to have a relatively broader range of ACE2 usage. On the other hand, the node at the MRCA of RaTG13 and MRCA1 (“MRCA2” in **Figure 4A**), was estimated to has limited ACE2 usage range. These data suggest that the amino acid substitution(s) between MRCA2 and MRCA1 conferred a broader ACE2 usage spectrum to sarbecovirus S. To identify the amino acid substitutions responsible for the expansion of ACE2 usage, we reconstructed and compared the amino acid sequences of the MRCA1 and MRCA2. We found that MRCA1 harbored seven substitutions when compared to MRCA2: K439N, K478T, Q483V, T484E, Y493Q, Y498H, and D501N (**Figure 4A**). Notably, all of these substitutions are located within the ACE2 binding interface of S (**Figure 4B**).

To experimentally assess the key substitution(s) that contribute to the expansion of the ACE2 usage range, the S expressing plasmids containing MRCA1 and MRCA2 were prepared. Using the plasmid expressing the S protein of Guangdong/1/2019 S (GD2019), which has the broadest range of ACE2 usage (**Figures 1B and S1**), as the backbone, we prepared two chimeric S expression plasmids, GD2019-MRCA1 and GD2019-MRCA2 (**Figure 4C**). As expected, the broadness scores of both GD2019-MRCA1 and parental GD2019 were significantly increased compared to GD2019-MRCA2 (*P* = 0.0060 and *P* = 0.0087) (**Figure 4D**). This result supports the hypothesis that either of these seven substitution(s) contributed to the expansion of the range of ACE2 use.

To determine the substitution(s) responsible for the broader ACE2 usage range of the SC2r clade, we prepared a series of derivatives bearing either or all of the substitutions based on the S proteins of GD2019-MRCA1, GD2019-MRCA2, and the parental GD2019. In the case of GD2019-MRCA2 derivatives, as expected, the derivative with all seven substitutions [“All (7sub)” in **Figure 4E**] gained the function of expanding the range of ACE2 use. In particular, we found that only the Y498H substitution significantly increased the broadness score (*P* < 0.0001) (**Figures 4E and S4A**). The co-structure model of SARS-CoV-2 S and human ACE2 (PDB:6M0J)^30^ showed that the region of ACE2 surrounding residue 498 in S is negatively charged, suggesting that this ACE2 region can lead to an electrostatic interaction between itself and the positively charged H498 (**Figure S3B**). These results suggest that Y498H is a key substitution for the broad spectrum of ACE2 use of SC2r clade.

In the experiments with the derivatives of GD2019-MRCA1 and the parental GD2019, which are designed to detect the substitution(s) that reduce the broadness score, the derivatives with all seven substitutions [“All (7sub)” in **Figures 4F and 4G**] lost the ability to utilize a broad range of ACE2 proteins. Particularly, the three substitutions, N439K, Q493Y, and N501D, reduced the broadness scores in both experiments (**Figures 4F and 4G**).

Since the experiment based on the GD2019-MRCA2 (**Figure 4E**) was conducted by single or all substitutions, it is possible to assume that combinational substitutions can affect the broad spectrum of ACE2 use. Considering the results of the derivatives of GD2019-MRCA1 and the parental GD2019 (**Figures 4F and 4G**), it is hypothesized that the combinational substitutions of the three residues positioned at 439, 493, and 501 can increase the broadness score of GD2019-MRCA1. To address this possibility, the GD2019-MRCA2 S derivatives with substitution at two or three residues of these positions were generated and pseudovirus assay was performed. As shown in **Figure 4H**, the combination of these three substitutions (K439N/Y493Q/D501N) increased the broadness score of GD2019-MRCA2. On the other hand, the three double substitution derivatives (K439N/Y493Q, K439N/D501N, and Y493Q/D501N) did not increase the broadness score (**Figures 4H and S4D**). These results suggest that, in addition to the Y498H substitution, the substitution of the three residues positioned at 439, 493 and 501 confer to increase the broad spectrum of ACE2 use of the SC2r clade.

To address how the three substitutions (K439N/Y493Q/D501N) increase the broad ACE2 usage, we compared the co-structure models of human ACE2 with MRCA2 and MRCA1. As shown in **Figure S3C**, N439 in MRCA1 formed hydrogen bond with P499, while K439 in MRCA2 did not. In addition, the position of Y493 in MRCA2 RBD was different from Q493 in MRCA1 because of the bulky side chain of Y493. These substitutions potentially affect the local structure of the residue 501 in RBD. D501 in MRCA2 formed a salt bridge with K353 in human ACE2, leaving the negative charged E38 residue in hydrophobic region. On the other hand, N501 of MRCA1 formed a hydrogen bond with Y41 of human ACE2, which led to forming salt bridge between E38 and K353 in human ACE2. These observations suggest that K439N, Y493Q and D501N expand the ACE2 usage range by epistasis. Altogether, these results suggest that (1) the Y498H substitution; and (2) the K439N, Y493Q, and D501N substitutions contribute to the expansion of the ACE2 usage range of the SC2r clade.

## Discussion

Numerous sarbecoviruses circulate in *Rhinolophus* bats primarily in East and Southeast Asia but also in Europe and Africa. The fact that SARS-CoV and SARS-CoV-2 have emerged in humans in the last two decades suggests that similar events may occur in the future. Understanding the determinants of sarbecovirus host range is critical for pandemic preparedness. Here, we used pseudoviruses expressing the S proteins of 53 sarbecoviruses and cells expressing ACE2 proteins of 17 mammalian species (including 12 *Rhinolophus* bat species) to elucidate the substitutions controlling the range of ACE2 usage for sarbecovirus infection.

We revealed that sarbecoviruses in the SC2r clade, including SARS-CoV-2, have a wider ACE2 usage range compared to other sarbecoviruses. By inferring the ACE2 usage range of ancestral sarbecoviruses we showed that the wider ACE2 usage range of SC2r clade was acquired during the evolution. We also found that the wider ACE2 usage range is determined by acquiring residues H498 and N439/Q493/N501 in their RBDs (**Figure 4**). Previous studies showed that SARS-CoV-2 and the variants has broad host range^35–38^. In addition, these studies reported that positions 417, 449, 486, 493, 498, 501 in RBM are contributed to the broad host range of SARS-CoV-2 and the variants^37,38^. Consistent with these studies, positions 493, 498 and 501 contributed to the expansion of ACE2 usage range. However, positions 417, 449 and 486 did not contribute to the expansion. Furthermore, the acquisition of N439 was found to contribute to the expansion through an epistatic effect with residues 493 and 501 (**Figure S3C**). Moreover, it is noteworthy that the acquisition of H498 was found to expand the ACE2 usage range independently. H498 is carried by SARS-CoV-2-related coronaviruses and some pangolin coronaviruses belonging to the Guangxi (GX) strain. The expansion of ACE2 usage range by Y498H substitution might be caused by the electrostatic interaction with the negatively charged region in ACE2 (**Figure S3B**). Our data suggest that ACE2 tropism can be substantially altered by as few as a single substitution in the S protein.

The S protein of MRCA1 contains H498, whereas that of SARS-CoV-2 contains Q498. A previous study showed that the H498Q substitution does not affect infectivity in the cells expressing human ACE2 but reduces infectivity in the cells expressing *Rhinolophus* bat ACE2s^39^. These observations suggest that H498 contributed to the expansion of sarbecovirus host range but is not critical for human spillover. Notably, the Y498H substitution allows the use of ACE2 proteins not only from various *Rhinolophus* bats but also from civets and raccoon dogs, which are presumed to have been involved as intermediate hosts for SARS-CoV^40^ and SARS-CoV-2^3,41,42^. In other words, it is reasonable to assume that the expansion of host range can potentially increase the risk of outbreaks in the human population.

As shown in **Figure 1**, BtKY72 belongs to Clade 3 and is phylogenetically distant from SARS-CoV-2. Additionally, the percentage similarity between the BtKY72 and SARS-CoV-2 S proteins is only 72.1% (73.2% in the RBD). Nevertheless, the use of either *R. ferrumequinum* or human ACE2 was determined by a single amino acid residue at site 493 (K or Q) of the S protein (**Figure 2**). These results suggest that viral spillover potential within sarbecoviruses is not limited to phylogenetic or genetic similarity and that only one or a few substitutions can drastically alter sarbecovirus host tropism.

Although RshSTT200 and Ra22QT77 belong to Clade 1b, these S proteins have the unique feature of not using human ACE2 for infection. Here, we revealed that four substitutions (T346R, P486F, D487N, and Y496G) conferred the ability to use human ACE2. In the case of RshSTT200 S, the use of human ACE2 or the ACE2 of its natural host (i.e., *R. shameli*), as determined by the four substitutions above, was mutually exclusive. Additionally, a similar result was observed for the S protein of BtKY72, a Clade 3 virus, where residue 493 (K or Q) determines the availability of human or *R. ferrumequinum* ACE2 usage in a mutually exclusive manner. However, it was of interest that the four substitutions in the Ra22QT77 S conferred the ability to use both the ACE2 of its natural host (i.e., *R. affinis*) and human ACE2. These results suggest that sarbecoviruses can expand their host range by acquiring certain substitutions in the S protein.

Cryo-EM structural analysis revealed that residue 38 of *R. cornutus* ACE2 is modified by N-linked glycosylation and interacts with N494 of Rc-o319 S (**Figure 3**). Pseudovirus infectivity assay showed that the loss of the N38-linked glycan of *R. cornutus* ACE2 (the N38D or T40A substitution) increased the infectivity of Rc-o319, suggesting that the N38-linked glycan of *R. cornutus* ACE2 inhibits the binding of Rc-o319 S. This is reminiscent of the finding that the N90-linked glycan of human ACE2 reduced the binding affinity with the S protein of the ancestral SARS-CoV-2^43,44^. In contrast to the N38-linked glycan, we demonstrated the importance of Y41 in *R. cornutus* ACE2 for interacting with Rc-o319 S. Since the Y498A substitution of Rc-o319 S abolished infectivity, our results suggest that the interaction between Y498 of Rc-o319 S and Y41 of *R. cornutus* ACE2 is essential for their binding. Previous studies showed that Y41 of human ACE2 and residue H498 of SARS-CoV-2 S mutant possessing Q498H form a π-π interaction between the aromatic rings in both residues^45,46^. Therefore, the binding of Rc-o319 S with *R. cornutus* ACE2 could be determined by this π-π interaction.

In summary, we demonstrated the mechanisms pertaining the sarbecovirus host tropism evolution in their natural hosts (*Rhinolophus* bats and pangolins), potential intermediate hosts (civets and raccoon dogs), and humans. We showed that the ACE2 tropism of sarbecoviruses can be easily altered and the spectrum of ACE2 usage can be broadened by only a few substitutions in the S protein. The expansion of host range in these viruses can increase the potential risk of viral spillover events that may eventually lead to disease outbreaks in humans. Our results highlight not only the importance of collecting novel viral sequences but also that of experimentally elucidating the phenotypes of viral proteins, such as the receptor tropism of S proteins. Monitoring sarbecoviruses circulating in the wild and characterizing the range of ACE2 usage are critical to preparing for a future pandemic.

## Supporting information

Key Resources Table

PDB Verification Report

## Acknowledgments

We gratefully acknowledge all data contributors, i.e. the Authors and their Originating laboratories responsible for obtaining the specimens, and their Submitting laboratories for generating the genetic sequence and metadata and sharing via the GISAID Initiative, on which this research is based. We would like to express our gratitude to all the members of Division of Systems Virology, The Institute of Medical Science, The University of Tokyo. We thank Dr. Shin Murakami (The University of Tokyo) and Dr. Kenzo Tokunaga (National Institute of Infectious Diseases, Japan) for sharing materials.

This study was supported in part by AMED ASPIRE Program (24jf0126002, to Kei Sato); AMED SCARDA Japan Initiative for World-leading Vaccine Research and Development Centers “UTOPIA” (243fa627001h0003, to Kei Sato); AMED SCARDA Program on R&D of new generation vaccine including new modality application (243fa727002, to Kei Sato); AMED Research Program on Emerging and Re-emerging Infectious Diseases (fk0108583, fk0108690, to Kei Sato); AMED Japan Program for Infectious Diseases Research and Infrastructure (Collaborative Research via Overseas Research Centers) (24wm0225041, to Kei Sato); JST PRESTO (JPMJPR22R1, to Jumpei Ito); JSPS KAKENHI Fund for the Promotion of Joint International Research (International Leading Research) (JP23K20041, to Kei Sato); JSPS KAKENHI Grant-in-Aid for Early-Career Scientists (JP23K14526, to Jumpei Ito); JSPS Research Fellow DC1 (23KJ0710, to Yusuke Kosugi); JSPS KAKENHI Grant-in-Aid for Scientific Research A (JP24H00607, to Kei Sato); Mitsubishi UFJ Financial Group, Inc. Vaccine Development Grant (to Jumpei Ito and Kei Sato); grants from the Japanese Government Ministry of Education, Culture, Sports, Science and Technology Scholarship–Research Category (220235, to Jarel Elgin Tolentino; 240042, Maximilian Stanley Yo); the Platform Project for Supporting Drug Discovery and Life Science Research (Basis for Supporting Innovative Drug Discovery and Life Science Research (BINDS)) from AMED (JP24ama121002 (support number 3272, to Osamu Nureki) and JP24ama121012 (supporting number S02820001 and S02820002, to Kei Sato)); Shionogi Infectious Disease Research Promotion Foundation (to Wataru Shihoya).

## STAR⍰METHODS

### KEY RESOURCES TABLE

### RESOURCE AVAILABILITY

#### Lead Contact

Further information and requests for resources and reagents should be directed to and will be fulfilled by the Lead Contact, Kei Sato.

#### Materials Availability

All unique reagents generated in this study are listed in the Key Resources Table and available from the Lead Contact with a completed Materials Transfer Agreement.

#### Data and code availability

All databases/datasets used in this study are available from the GenBank (https://www.ncbi.nlm.nih.gov/genbank/) and GISAID databases (https://www.gisaid.org; EPI_ISL_410721; EPI_ISL_412977; EPI_ISL_852604; EPI_ISL_852605). Code and data used in this study are available on the following GitHub repository: https://github.com/TheSatoLab/ACE2_tropism. The atomic coordinates and cryo-EM maps of the structures of the Rc-o319 RBD/*R. cornutus* ACE2 complex (PDB:9M3F, EMD:63603) generated in this study have been deposited in the Protein Data Bank (https://www.rcsb.org/), and Electron Microscopy Data Bank (https://www.ebi.ac.uk/emdb/). Any additional information required to reanalyze the data reported in this work is available from the lead contact upon request.

#### Nucleotide sequence data collection

All 250 whole genome sequences of known sarbecoviruses used in Pekar *et al.* (2023)^47^ in addition to a single civet-infecting SARS-CoV genome (civet007, NCBI accession: AY572034) were retrieved. Also, the nucleotide sequences of ACE2 of various species were collected from NCBI GenBank (download date, 18 December 2023). Information on the sarbecoviruses and ACE2 sequences are summarized in **Table S1** and **Table S2.**

#### Molecular phylogenetic analysis

To assess the phylogenetic relatedness between all sarbecovirus RBDs, we first extracted the RBD-encoding part of the 251 genomes (using the SARS-CoV-2 RBD as a reference, i.e., amino positions 319-541 in the SARS-CoV-2 Wuhan-Hu-1 strain S protein) and aligned the coding sequences on the nucleotide level using the localpair option of MAFFT v7.505^48^. Three additional RBDs which were nearly identical to pangolin sarbecovirus MP789, but had few RBD amino acid differences were also included in the analysis (Guangdong/1/2019: EPI_ISL_410721, cDNA8: MT799521.1, cDNA16: MT799523.1). We inferred a maximum likelihood (ML) tree based on this alignment using IQ-TREE2 version 2.3.4^49^ under a GTR+F+I+R4 substitution model. Node support was assessed with 10,000 Ultrafast bootstrap replicates^50^. Based on this tree (**Figure S1**), we manually selected 53 representative sarbecovirus RBDs covering all diversity in the tree. We extracted the RBD sequences for these 53 representatives and constructed an amino acid alignment using MUSCLE v3.8.1551^51^ with default options. An ML tree was then reconstructed for this amino acid alignment using IQ-TREE2 under the best substitution model selected automatically by ModelFinder^52^ with 1000 bootstrap replicates (Figure 1A). This ML tree, rooted by the Clade 3 sarbecoviruses was used to perform ancestral reconstruction of the internal node RBD amino acid sequences (Figure 4A), using TreeTime v0.11.3^53^.

In Figure 1B, the ML tree of amino acid sequences of *Rhinolophus* ACE2 and other five mammalian ACE2 (raccoon dog, raccoon, civet, pangolin and human) was constructed by the following procedures. An MSA of ACE2 proteins was constructed using MUSCLE v3.8.1551^51^ implemented in MEGA10 v10.1.7^54^. The ML tree was reconstructed using MEGA10 under the GTR+G substitution model with 100 bootstrap replicates.

#### Cell culture

HEK293 cells (a human embryonic kidney cell line; ATCC, CRL-1573), LentiX-293T cell line (a human embryonic kidney cell line; ATCC, Takara, Cat# 632180) and HOS-ACE2/TMPRSS2 cells, HOS cells (a human osteosarcoma cell line; ATCC CRL-1543) stably expressing human ACE2 and TMPRSS2^55,56^ were maintained in Dulbecco’s modified Eagle’s medium (high glucose) (Sigma-Aldrich, Cat# 6429-500ML) containing 10% fetal bovine serum and 1% penicillin-streptomycin (Sigma-Aldrich, Cat# P4333-100ML). HEK293 GnTI^-^ cells (ATCC, CRL-3022) were cultured in FreeStyle™ 293 Expression Medium (Gibco, Cat# 12338018) supplemented with 2% FBS (Nichirei, Cat# 175012) with 130 rpm under 8% CO2 atmosphere at 37°C. The HOS-TMPRSS2 cells that stably express *Rhinolophus* bat ACE2 were generated as described below (see “Generation of HOS-TMPRSS2 cells stably expressing a variety of ACE2 proteins” section) and maintained in Dulbecco’s modified Eagle’s medium (high glucose) (Sigma-Aldrich, Cat# 6429-500ML) containing 10% fetal bovine serum and 1% penicillin-streptomycin (Sigma-Aldrich, Cat# P4333-100ML), zeocin (50 μg/mL; InvivoGen, Cat#ant-zn-1) and G418 (400 μg/ml; Nacalai Tesque, Cat# G8168-10ML).

#### Plasmid construction

A part of plasmids expressing the human codon-optimized S proteins of sarbecoviruses were prepared in our previous study^25^. or the ACE2 proteins Plasmids expressing the human codon-optimized S proteins of sarbecoviruses (summarized in **Table S1**) were synthesized by a gene synthesis service (Fasmac). Plasmids expressing the ACE2 proteins summarized in **Table S2** were also synthesized by a gene synthesis service (Fasmac). Plasmids expressing the derivatives of codon-optimized S proteins of sarbecoviruses and *Rhinolophus* bat ACE2 were generated by site-directed overlap extension PCR using the primers summarized in **Table S3.**

The resulting PCR fragment was cloned into the KpnI/NotI site of backbone pCAGGS vector^57^ (for the S expression plasmids) or the BamHI/MluI site of pWPI-ACE2-zeo (for ACE2 expression plasmids)^56^ with 3×FLAG-tag at the C-terminus using In-Fusion® HD Cloning Kit (Takara, Cat# Z9650N). Nucleotide sequences were determined by DNA sequencing services (Eurofins), and the sequence data were analyzed by SnapGene v8.0.1 (SnapGene software).

#### Generation of HOS-TMPRSS2 cells stably expressing a variety of ACE2 proteins

HOS-TMPRSS2 cells stably expressing ACE2 proteins of *R. affinis*, *R. cornutus*, *R. sinicus*, *R. ferrumequinum*, *R. shameli*, *R. pearsonii*, *R. pusillus*, *R. macrotis*, human, and pangolin were prepared in our previous study ^25^. To prepare lentiviral vectors expressing ACE2, LentiX-293T cells (500,000 cells) were cotransfected with 0.9 μg of psPAX2-IN/HiBiT, 0.2 μg of pCMV-VSV-G-RSV-Rev, and 0.9 μg of pWPI-ACE2-zeo using TransIT-293 (Takara, Cat# MIR2704) according to the manufacturer’s protocol. After 48 hours of transfection, the supernatant including lentivector particles was collected. HOS-TMPRSS2 cells (100,000 cells) were then transduced with the ACE2-expressing lentiviral vector. After 48 hours post transduction, transduced cells were maintained for zeocin (50 μg/mL; Invivogen, Cat# ant-zn-1) and G418; 400 μg/mL; Nacarai Tesque, Cat# 09380-44; selections for 14 days.

#### Pseudovirus assay

Pseudovirus assay was performed as previously described^58–62^. Briefly, HIV-1-based, luciferase-expressing reporter viruses were pseudotyped with the S proteins of sarbecoviruses and their derivatives. Lenti-293T cells (500,000 cells) were cotransfected with 0.8 μg psPAX2-IN/HiBiT^55^, 0.8 μg pWPI-Luc2^63^, and 0.4 μg plasmids expressing parental S or its derivatives using TransIT-293 (Takara, Cat# MIR2704) according to the manufacturer’s protocol. Two days posttransfection, the culture supernatants were harvested, and the pseudoviruses were stored at –80°C until use. For pseudovirus infection, the amount of input virus was normalized to the HiBiT value measured by NanoGlo HiBiT lytic detection system (Promega, Cat# N3040) as previously described^63^. In this system, HiBiT peptide is produced with HIV-1 integrase and forms NanoLuc luciferase with LgBiT, which is supplemented with substrates. In each pseudovirus particle, the detected HiBiT value is correlated with the amount of the pseudovirus capsid protein, HIV-1 p24 protein^63^. Therefore, we calculated the amount of HIV-1 p24 capsid protein based on the HiBiT value measured, according to the previous paper^63^. The amount of HIV-1 p24 antigen used in the assay was 4 ng (Figures 1B**, 2A, 2G, 2H, 2K and 4D-4G**), 5 ng (Figures 2C **and 2D**), 40 ng (Figures 2F **and 2I**) or 10 ng (Figures 3B **and 3C**). For target cells, the HOS-TMPRSS2 cells stably expressing a variety of *Rhinolophus* bat ACE2 (Figures 1B**, 2A, 2F-2J and 4D-4G**) and the HEK293 cells transfected with the plasmids expressing *R. cornutus*, *R. ferrumequinum* and human ACE2 and their derivatives with PEI-Max (Polysciences, Cat# 24765-1) (Figures 2C**, 2D, 3B and 3C**) were used. Two days postinfection, the infected cells were lysed with a Bright-Glo Luciferase Assay System (Promega, cat# E2620) and the luminescent signal was measured using a GloMax Explorer Multimode Microplate Reader (Promega).

#### Inferring change in ACE2 usage range within the sarbecovirus lineage

We inferred change in ACE2 usage range within the sarbecovirus lineage using the phylogenetic tree reconstructed from sarbecovirus RBDs (Figure 1A) and infectivity data obtained from the pseudovirus infection assay (Figure 1B). First, we calculated the standardized Z score for the infectivity of sarbecoviruses in HOS-TMPRSS2 cells expressing ACE2 from various host species. The broadness score was calculated as the median of the standardized infectivity and used to represent the ACE2 usage range for each sarbecovirus. Next, we estimated the ACE2 usage ranges of sarbecovirus ancestral nodes in the phylogenetic tree using the fastAnc function in phytools R package v2.1-1^64^, which reconstructs ancestral states for continuous traits under the maximum likelihood model. We then compared the medians of standardized infectivity between ancestral nodes and their adjacent descendant nodes to infer changes in the ACE2 usage range. All analyses were performed in R v4.4.0 (https://www.R-project.org/).

#### Recombinant protein for structural analysis

The Rc-o319 Ectodomain and *R. cornutus* ACE2 sequences, each containing a C-terminal 6×His-tag, were independently subcloned into pHL-sec vectors (Addgene, Cat# 99845)^65^. Both proteins were co-expressed and secreted by HEK293 GnTI^-^ cells. The cells were removed after 5 days by centrifuging at 5,000 x g for 10 min, and the supernatant containing the secreted material was mixed with 20 mM Tris (pH 7.5) and 150 mM NaCl. The supernatant was incubation with Ni Sepharose™ excel resin (Cytiva, Cat# 17371201) at 4 °C for an hour. The recovered resin was further washed with 10 column volumes of buffer containing 20 mM Tris (pH 7.5), and 150 mM NaCl, 20 mM imidazole. Proteins were then eluted with 20 mM Tris (pH 7.5), 150 mM NaCl, and 400 mM imidazole. The eluted fractions were concentrated and loaded onto a Superdex™ 200 Increase 10/300 GL size exclusion column (Cytiva, Cat# 28990944) equilibrated with buffer containing 20 mM Tris (pH 7.5) and 150 mM NaCl.

#### Cryo-EM sample preparation and data collection

These pacificated proteins were concentrated and added n-Octyl-β-D-glucoside (OG) (Nacarai Tesque, Cat# 25535-82) to a final concentration of 0.01%. The purified and concentrated complexes (8.3 mg/mL, 3 µL) were applied on the glow-discharged Quantifoil Au grids (R1.2/1.3, 300 mesh) (Quantifoil Micro Tools GmbH) and plunged into the liquid ethane using a Vitrobot Mark IV (FEI). Micrographs were collected using a 300 kV-Titan Krios G3i microscope (Thermo Fisher Scientific) equipped with a BioQuantum K3 imaging filter and a K3 direct electron detector (Gatan). In total, 14,895 movies were acquired using EPU software with a calibrated pixel size of 0.83 Å/pix and a defocus range of −0.8 to −1.6 μm.

#### Cryo-EM image processing, model building, and model refinement

Data processing was performed using cryoSPARC v4.4.036^66^. Dose-fractionated movies were aligned using Patch motion correction and contrast transfer functions (CTF) were estimated using Patch CTF Estimation. Detailed data processing is shown in Figure S4. The density map was sharpened using DeepEMhancer version 0.14^67^. The quality of the density maps was sufficient to build the model manually using COOT for Windows version 0.9.8.93 ^68,69^. Model building was manually performed based on the BANAL-20-236 complex structure (PDB:8HXK)^39^. The model was then refined using phenix.real_space_refine version 1.19^70,71^.

#### Protein structure model

In Figure 2D, 5 models of the structure of *R. ferrumequinum* ACE2 were estimated using AlphaFold3^72^. Evaluation of the models were performed using pLDDT scores, pTM scores and ipTM scores^73^ then the best model for the co-structure was selected. The crystal co-structure of BtKY72 S RBD and *R. landeri* ACE2 (PDB:8K4U)^31^ were used. To infer interaction between S RBD and ACE2, the structure of *R. landeri* ACE2 was replaced by the AlphaFold3 models of *R. ferrumequinum* ACE2. In **Figure S3C,** 5 models of the structural model of complex of MRCA1 or MRCA2 RBD and the human ACE2 were also estimated using AlphaFold3 in the same way. All protein structural analyses were performed using the PyMOL molecular graphics system v3.0.0 (Schrödinger) and Chimera X version 1.6^74^.

