## Supplementary material for "Molecular basis of sarbecovirus evolution and receptor tropism in natural hosts, potential intermediate hosts, and humans": Key Resources Table

| REAGENT or RESOURCE | SOURCE | IDENTIFIER |
| --- | --- | --- |
| Chemicals, peptides, and recombinant proteins | | |
| PEI-Max | Polysciences | Cat# 24765-1 |
| TransIT-293 transfection reagent | Mirus | Cat# MIR2704 |
| Fetal bovine serum | Sigma-Aldrich | Cat# 172012-500ML |
| Fetal bovine serum | Nichirei | Cat# 175012-500ML |
| Penicillin-streptomycin | Sigma-Aldrich | Cat# P4333-100ML |
| DMEM (high glucose) | Sigma-Aldrich | Cat# 6429-500ML |
| DMEM (high glucose) | Nacalai Tesque | Cat# 08458-16 |
| FreeStyle™ 293 Expression Medium | Gibco | Cat# 12338018 |
| zeocin | InvivoGen | Cat# ant-zn-1 |
| G418 | Nacalai Tesque | Cat# G8168-10ML |
| KpnI | New England Biolab | Cat# R3142S |
| NotI | New England Biolab | Cat# R1089S |
| BamHI | New England Biolab | Cat# R3136S |
| MluI | New England Biolab | Cat# R3198S |
| In-Fusion® HD Cloning Kit | Takara | Cat# Z9650N |
| PBS | Nacalai Tesque | Cat# 09154-85 |
| n-Octyl-β-D-glucoside | Nacalai Tesque | Cat# 25535-82 |
| Critical commercial assays | | |
| Nano Glo HiBiT lytic detection system | Promega | Cat# N3040 |
| Bright-Glo luciferase assay system | Promega | Cat# E2650 |
| Deposited data | | |
| Viral genome sequencing data of working viral stocks (see also **Table S4**) | This study | SRA: PRJDB14324  (https://www.ncbi.nlm.nih.gov/sra) |
| Cryo-EM map of the complex structure of Rc-o319 RBD/*R. cornutus* ACE2 | This study | EMD-xxx |
| Cryo-EM complex structure of Rc-o319 RBD/*R. cornutus* ACE2 | This study | PDB: xxx |
| Crystal complex structure of SARS-CoV-2 RBD/human ACE2 | Lan et al.^1^ | PDB:6M0J |
| Cryo-EM complex structure of BtKY72 RBD/*R. landeri* ACE2 | Su et al.^2^ | PDB:8K4U |
| Crystal complex structure of RshSTT182/200 RBD/human ACE2 | Hu et al.^3^ | PDB:7XBH |
| Cryo-EM complex structure of BANAL-20-236 RBD/human ACE2 | Ou et al.^4^ | PDB:8HXK |
| Experimental models: Cell lines | | |
| Human: HEK293 cells | ATCC | CRL-1573 |
| LentiX-293T | TaKaRa | Cat# 632180 |
| HEK293 GnTI^-^ cells | ATCC | CRL-3022 |
| Human: HOS-ACE2/TMPRSS2 cells | Ferreira et al. ^5^ and Ozono et al.^5,6^ | N/A |
| Human: HOS-TMPRSS2 cells | Fujita et al.^7^ | N/A |
| Oligonucleotides | | |
| Primers for the construction of plasmids expressing the codon-optimized S proteins of BA.2-bearing variants, see **Table S3** | This study | N/A |
| Recombinant DNA | | |
| Plasmid: pCAGGS | Niwa et al.^8^ | N/A |
| Plasmid: psPAX2-IN/HiBiT | Ozono et al.^9^ | N/A |
| Plasmid: pWPI-Luc2 | Ozono et al.^9^ | N/A |
| Plasmid: pWPI-MCS-Zeo | Fujita et al.^7^ | N/A |
| Plasmid: pHL-sec | Addgene | Cat# 99845 |
| Plasmid: pC-SARS-CoV-2 S | Ozono et al.^6^ | N/A |
| Plasmid: pC-BtKY72 S | This study | N/A |
| Plasmid: pC-RhGB01 S | This study | N/A |
| Plasmid: pC-BM48-31 S | This study | N/A |
| Plasmid: pC-RsSHC014 S | This study | N/A |
| Plasmid: pC-YN2020H S | This study | N/A |
| Plasmid: pC-BtRS-BetaCoV/YN2020-Q325 S | This study | N/A |
| Plasmid: pC-SARS-CoV S | This study | N/A |
| Plasmid: pC-Civet007 S | This study | N/A |
| Plasmid: pC-Rs7327 S | This study | N/A |
| Plasmid: pC-RsYN2016A S | This study | N/A |
| Plasmid: pC-LYRa11 S | This study | N/A |
| Plasmid: pC-WIV1 S | This study | N/A |
| Plasmid: pC-WIV16 S | This study | N/A |
| Plasmid: pC-Rc-os20 S | This study | N/A |
| Plasmid: pC-Rc-o319 S | This study | N/A |
| Plasmid: pC-Rc-kw8 S | This study | N/A |
| Plasmid: pC-Rc-mk2 S | This study | N/A |
| Plasmid: pC-MpCoV-GX S | This study | N/A |
| Plasmid: pC-GX-P5E S | This study | N/A |
| Plasmid: pC-GX-P5L S | This study | N/A |
| Plasmid: pC-GX-P4L S | This study | N/A |
| Plasmid: pC-GX-P2V S | This study | N/A |
| Plasmid: pC-RaTG13 S | This study | N/A |
| Plasmid: pC-Ra22QT77 S | This study | N/A |
| Plasmid: pC-RshSTT200 S | This study | N/A |
| Plasmid: pC-MpCoV-GD S | This study | N/A |
| Plasmid: pC-cDNA16 S | This study | N/A |
| Plasmid: pC-cDNA8 S | This study | N/A |
| Plasmid: pC-Guangdong/1/2019 S | This study | N/A |
| Plasmid: pC-BANAL-20-52 S | Fujita et al.^7^ | N/A |
| Plasmid: pC-BANAL-20-236 S | Fujita et al.^7^ | N/A |
| Plasmid: pC-Rp22DB159 S | This study | N/A |
| Plasmid: pC-CX1 S | This study | N/A |
| Plasmid: pC-RsYN04 S | This study | N/A |
| Plasmid: pC-Ra7909 S | This study | N/A |
| Plasmid: pC-GD2017H S | This study | N/A |
| Plasmid: pC-HKU3-9 S | This study | N/A |
| Plasmid: pC-YN2020A S | This study | N/A |
| Plasmid: pC-GD2017F S | This study | N/A |
| Plasmid: pC-Rm1 S | This study | N/A |
| Plasmid: pC Rs4081 S | This study | N/A |
| Plasmid: pC-JX2021M S | This study | N/A |
| Plasmid: pC-RmYN07 S | This study | N/A |
| Plasmid: pC-RsYN09 S | This study | N/A |
| Plasmid: pC-Anlong-103 S | This study | N/A |
| Plasmid: pC-RmYN02 S | This study | N/A |
| Plasmid: pC-RacCS203 S | This study | N/A |
| Plasmid: pC-GX2019A S | This study | N/A |
| Plasmid: pC-LN2020G S | This study | N/A |
| Plasmid: pC-BtRf-BetaCoV-HeB2013 S | This study | N/A |
| Plasmid: pC-RpYN06 S | This study | N/A |
| Plasmid: pC-bat-SL-CoVZC45 S | This study | N/A |
| Plasmid: pC-BtKY72 S K493Q | This study | N/A |
| Plasmid: pC-BtKY72 S T498Q | This study | N/A |
| Plasmid: pC-BtKY72 S K493Q/T498Q | This study | N/A |
| Plasmid: pC-RshSTT200 S T346R | This study | N/A |
| Plasmid: pC-RshSTT200 S P486F | This study | N/A |
| Plasmid: pC-RshSTT200 S D487N | This study | N/A |
| Plasmid: pC-RshSTT200 S Y496G | This study | N/A |
| Plasmid: pC-RshSTT200 S 4mut | This study | N/A |
| Plasmid: pC-RshSTT200 S ins | This study | N/A |
| Plasmid: pC-RshSTT200 S ins T346R | This study | N/A |
| Plasmid: pC-RshSTT200 S ins P486F | This study | N/A |
| Plasmid: pC-RshSTT200 S ins D487N | This study | N/A |
| Plasmid: pC-RshSTT200 S ins 4mut | This study | N/A |
| Plasmid: pC-SARS-CoV-2 S R346T | This study | N/A |
| Plasmid: pC-SARS-CoV-2 S F486P | This study | N/A |
| Plasmid: pC-SARS-CoV-2 S N487D | This study | N/A |
| Plasmid: pC-SARS-CoV-2 S G496Y | This study | N/A |
| Plasmid: pC-SARS-CoV-2 S 4mut | This study | N/A |
| Plasmid: pC-SARS-CoV-2 S del | This study | N/A |
| Plasmid: pC-SARS-CoV-2 S del R346T | This study | N/A |
| Plasmid: pC-SARS-CoV-2 S del F486P | This study | N/A |
| Plasmid: pC-SARS-CoV-2 S del N487D | This study | N/A |
| Plasmid: pC-SARS-CoV-2 del G496Y | This study | N/A |
| Plasmid: pC-SARS-CoV-2 S del 4mut | This study | N/A |
| Plasmid: pC-Ra22QT77 S T346R | This study | N/A |
| Plasmid: pC-Ra22QT77 S P486F | This study | N/A |
| Plasmid: pC-Ra22QT77 S D487N | This study | N/A |
| Plasmid: pC-Ra22QT77 S Y496G | This study | N/A |
| Plasmid: pC-Ra22QT77 S 4mut | This study | N/A |
| Plasmid: pC-Ra22QT77 S ins | This study | N/A |
| Plasmid: pC-Ra22QT77 S ins 4mut | This study | N/A |
| Plasmid: pC-Rc-o319 S Y498A | This study | N/A |
| Plasmid: pC-Rc-o319 S S500A | This study | N/A |
| Plasmid: pC-Rc-o319 S Y498A/S500A | This study | N/A |
| Plasmid: pC-Guangdong/1/2019 S N439K | This study | N/A |
| Plasmid: pC-Guangdong/1/2019 S T478K | This study | N/A |
| Plasmid: pC-Guangdong/1/2019 S V483Q | This study | N/A |
| Plasmid: pC-Guangdong/1/2019 S E484T | This study | N/A |
| Plasmid: pC-Guangdong/1/2019 S Q493Y | This study | N/A |
| Plasmid: pC-Guangdong/1/2019 S H498Y | This study | N/A |
| Plasmid: pC-Guangdong/1/2019 S N501D | This study | N/A |
| Plasmid: pC-Guangdong/1/2019 S 7mut | This study | N/A |
| Plasmid: pC-GD2019-CA1 S | This study | N/A |
| Plasmid: pC-GD2019-CA1 S N439K | This study | N/A |
| Plasmid: pC-GD2019-CA1 S T478K | This study | N/A |
| Plasmid: pC-GD2019-CA1 S V483Q | This study | N/A |
| Plasmid: pC-GD2019-CA1 S E484T | This study | N/A |
| Plasmid: pC-GD2019-CA1 S Q493Y | This study | N/A |
| Plasmid: pC-GD2019-CA1 S H498Y | This study | N/A |
| Plasmid: pC-GD2019-CA1 S N501D | This study | N/A |
| Plasmid: pC-GD2019-CA1 S 7mut | This study | N/A |
| Plasmid: pC-GD2019-CA2 S | This study | N/A |
| Plasmid: pC-GD2019-CA2 S K439N | This study | N/A |
| Plasmid: pC-GD2019-CA2 S K478T | This study | N/A |
| Plasmid: pC-GD2019-CA2 S Q483V | This study | N/A |
| Plasmid: pC-GD2019-CA2 S T484E | This study | N/A |
| Plasmid: pC-GD2019-CA2 S Y493Q | This study | N/A |
| Plasmid: pC-GD2019-CA2 S Y498H | This study | N/A |
| Plasmid: pC-GD2019-CA2 S D501N | This study | N/A |
| Plasmid: pC-GD2019-CA2 S 7mut | This study | N/A |
| Plasmid: pC-GD2019-CA2 S K439N/Y493Q | This study | N/A |
| Plasmid: pC-GD2019-CA2 S K439N/D501N | This study | N/A |
| Plasmid: pC-GD2019-CA2 S Y493Q/D501N | This study | N/A |
| Plasmid: pC-GD2019-CA2 S K439N/Y493Q/D501N | This study | N/A |
| Plasmid: pWPI-*R. cornutus* ACE2-3xFLAG-zeo | This study | N/A |
| Plasmid: pWPI-*R. pusillus* ACE2-3xFLAG-zeo | This study | N/A |
| Plasmid: pWPI-*R. macrotis* ACE2-3xFLAG-zeo | Fujita et al.^7^ | N/A |
| Plasmid: pWPI-*R. sinicus* ACE2-3xFLAG-zeo | Fujita et al.^7^ | N/A |
| Plasmid: pWPI-*R. thomasi* ACE2-3xFLAG-zeo | This study | N/A |
| Plasmid: pWPI-*R. shameli* ACE2-3xFLAG-zeo | Fujita et al.^7^ | N/A |
| Plasmid: pWPI-*R. affinis* ACE2-3xFLAG-zeo | Fujita et al.^7^ | N/A |
| Plasmid: pWPI-*R. pearsonii* ACE2-3xFLAG-zeo | Fujita et al.^7^ | N/A |
| Plasmid: pWPI-*R. alcyone* ACE2-3xFLAG-zeo | This study | N/A |
| Plasmid: pWPI-*R. landeri* ACE2-3xFLAG-zeo | This study | N/A |
| Plasmid: pWPI-*R. sp.* ACE2-3xFLAG-zeo | This study | N/A |
| Plasmid: pWPI-*R. ferrumequinum* ACE2-3xFLAG-zeo | Fujita et al.^7^ | N/A |
| Plasmid: pWPI-raccoon dog ACE2-3xFLAG-zeo | This study | N/A |
| Plasmid: pWPI-raccoon ACE2-3xFLAG-zeo | This study | N/A |
| Plasmid: pWPI-civet ACE2-3xFLAG-zeo | This study | N/A |
| Plasmid: pWPI-pangolin ACE2-3xFLAG-zeo | Fujita et al.^7^ | N/A |
| Plasmid: pWPI-human ACE2-3xFLAG-zeo | Fujita et al.^7^ | N/A |
| Plasmid: pWPI-*R. ferrumequinum* ACE2 D31K-3xFLAG-zeo | This study | N/A |
| Plasmid: pWPI-*R. ferrumequinum* ACE2 H41Y-3xFLAG-zeo | This study | N/A |
| Plasmid: pWPI-*R. ferrumequinum* ACE2 D31K/H41Y-3xFLAG-zeo | This study | N/A |
| Plasmid: pWPI-human ACE2 K31D-3xFLAG-zeo | This study | N/A |
| Plasmid: pWPI-human ACE2 Y41H-3xFLAG-zeo | This study | N/A |
| Plasmid: pWPI-human ACE2 K31D/Y41H-3xFLAG-zeo | This study | N/A |
| Plasmid: pWPI-*R. cornutus* ACE2 K27A-3xFLAG-zeo | This study | N/A |
| Plasmid: pWPI-*R. cornutus* ACE2 D31A-3xFLAG-zeo | This study | N/A |
| Plasmid: pWPI-*R. cornutus* ACE2 N38D-3xFLAG-zeo | This study | N/A |
| Plasmid: pWPI-*R. cornutus* ACE2 T40A-3xFLAG-zeo | This study | N/A |
| Plasmid: pWPI-*R. cornutus* ACE2 Y41A-3xFLAG-zeo | This study | N/A |
| Plasmid: pWPI-*R. cornutus* ACE2 Q42A-3xFLAG-zeo | This study | N/A |
| Plasmid: pWPI-*R. cornutus* ACE2 E75A-3xFLAG-zeo | This study | N/A |
| Plasmid: pWPI-*R. cornutus* ACE2 K353A-3xFLAG-zeo | This study | N/A |
| Plasmid: pWPI-*R. cornutus* ACE2 7mut-3xFLAG-zeo | This study | N/A |
| Plasmid: pHLsec-Rc-o319 S ectodomain-6xHis | This study | N/A |
| Plasmid: pHLsec-*R. cornutus* ACE2 (19-615)-6xHis | This study | N/A |
| Software and algorithms | | |
| MAFFT v7.505 | Katoh et al.^10^ | https://mafft.cbrc.jp/alignment/server/index.html |
| MUSCLE v3.8.1551 | Edgar^11^ | https://www.drive5.com/muscle/ |
| IQ-TREE multicore version 2.3.4 | Minh et al.^12^ | https://github.com/iqtree/iqtree2 |
| MEGA10 v10.1.7 | Kumar et al.^13^ | https://www.megasoftware.net/ |
| Treetime v0.11.3 | Sagulenko et al.^14^ | https://github.com/neherlab/treetime |
| SnapGene v8.0.1 | SnapGene software | https://www.snapgene.com/ |
| R v4.4.0 | The R Foundation | https://www.r-project.org/ |
| Phytools R package v2.1-1 | Reveil^15^ | https://github.com/liamrevell/phytools |
| EPU software | Thermo Fisher Scientific | https://www.thermofisher.com/jp/ja/home/about-us/events/industrial/smart-epu.html |
| cryoSPARC v4.4.036 | Punjani et al.^16^ | https://cryosparc.com/ |
| DeepEMhancer | Sanchez-Garcia et al.^17^ | https://github.com/rsanchezgarc/deepEMhancer |
| COOT for Windows version 0.9.8.93 | Emsley and Cowtan ^18,19^ | https://www2.mrc-lmb.cam.ac.uk/personal/pemsley/coot/ |
| phenix.real_space_refine | Adams et al. ^20^ and Afonine et al.^20,21^ | https://www.phenix-online.org/documentation/reference/real_space_refine.html |
| AlphaFold3 | Abramson et al.^22^ | https://golgi.sandbox.google.com/ |
| PyMOL molecular graphics system v3.0.0 | Schrödinger | https://www.python.org |
| Chimera X version 1.6 | Goddard et al. ^23^ | https://www.rbvi.ucsf.edu/chimerax/ |
| Other | | |
| NCBI GenBank | NCBI | https://www.ncbi.nlm.nih.gov/ |
| GloMax explorer multimode microplate reader 3500 | Promega | N/A |
| GISAID database | Khare et al.^24^ | https://www.gisaid.org/ |
| Quantifoil Au grids (R1.2/1.3, 300 mesh) | Quantifoil Micro Tools GmbH | N/A |
| 300 kV-Titan Krios G3i microscope | Thermo Fisher Scientific | N/A |
| BioQuantum K3 imaging filter | Gatan | N/A |
| K3 direct electron detector | Gatan | N/A |
