## Supplementary Information for "Molecular basis of sarbecovirus evolution and receptor tropism in natural hosts, potential intermediate hosts, and humans"

Supplementary Figures S1-S4
Supplementary Tables S1-S4

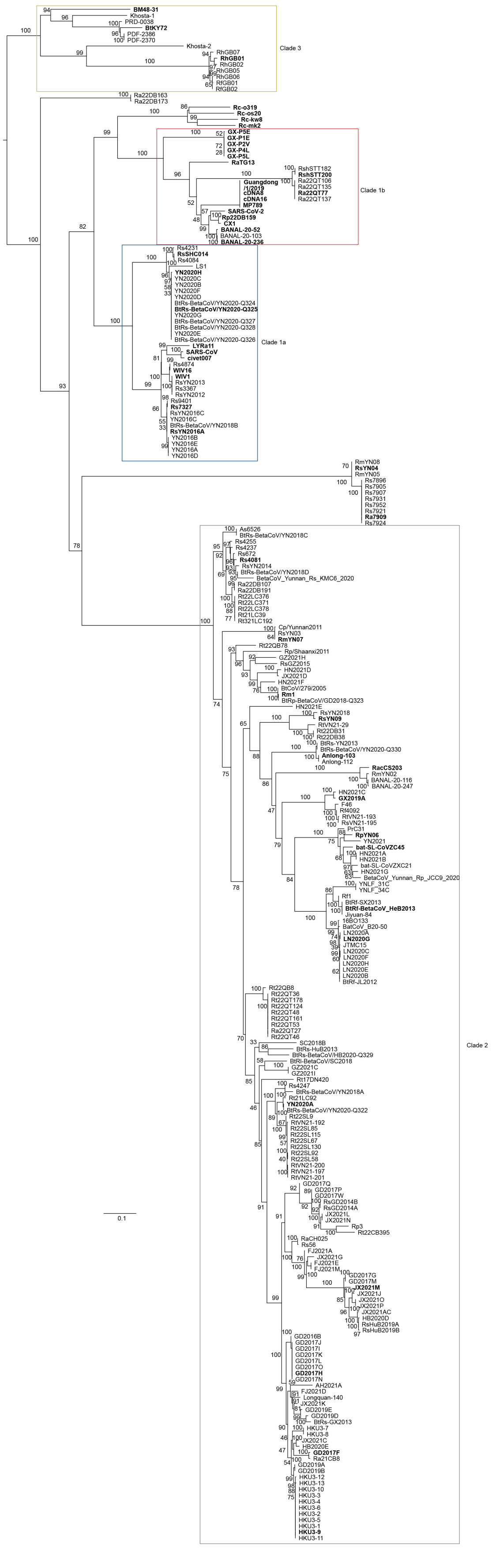

Clade 3
BM48-31
Khosta-1
PRD-0038
BtKY72
PDF-2386
PDF-2370
Khosta-2
RhGB07
RhGB01
RhGB02
RhGB05
RhGB06
RfGB01
RfGB02
Ra22DB163
Ra22DB173
Rc-o319
Rc-os20
Rc-kw8
Rc-mk2
Clade 1b
GX-P5E
GX-P1E
GX-P2V
GX-P4L
GX-P5L
RaTG13
RshSTT182
RshSTT200
Ra22QT106
Ra22QT135
Ra22QT77
Ra22QT137
Guangdong /1/2019
cDNA8
cDNA16
MP789
SARS-CoV-2
Rp22DB159
CX1
BANAL-20-52
BANAL-20-103
BANAL-20-236
Clade 1a
Rs4231
RsSHC014
Rs4084
LS1
YN2020H
YN2020C
YN2020B
YN2020F
YN2020D
BtRs-BetaCoV/YN2020-Q324
BtRs-BetaCoV/YN2020-Q325
YN2020G
BtRs-BetaCoV/YN2020-Q327
BtRs-BetaCoV/YN2020-Q328
YN2020E
BtRs-BetaCoV/YN2020-Q326
LYRa11
SARS-CoV
civet007
Rs4874
WIV16
WIV1
RsYN2013
Rs3367
RsYN2012
Rs9401
Rs7327
RsYN2016C
YN2016C
BtRs-BetaCoV/YN2018B
RsYN2016A
YN2016B
YN2016E
YN2016A
YN2016D
RmYN08
RsYN04
RmYN05
Rs7896
Rs7905
Rs7907
Rs7931
Rs7952
Rs7921
Ra7909
Rs7924
Clade 2
As6526
BtRs-BetaCoV/YN2018C
Rs4255
Rs4237
Rs672
Rs4081
RsYN2014
BtRs-BetaCoV/YN2018D
BetaCoV_Yunnan_Rs_KMC6_2020
Ra22DB107
Ra22DB191
Rt22LC376
Rt22LC371
Rt22LC378
Rt21LC39
Rt321LC192
Cp/Yunnan2011
RsYN03
RmYN07
Rt22QB78
Rp/Shaanxi2011
GZ2021H
RsGZ2015
HN2021D
JX2021D
HN2021F
BtCoV/279/2005
Rm1
BtRp-BetaCoV/GD2018-Q323
HN2021E
RsYN2018
RsYN09
RtVN21-29
Rt22DB31
Rt22DB38
BtRs-YN2013
BtRs-BetaCoV/YN2020-Q330
Anlong-103
Anlong-112
RacCS203
RmYN02
BANAL-20-116
BANAL-20-247
HN2021C
GX2019A
F46
Rf4092
RtVN21-193
RsVN21-195
PrC31
RpYN06
YN2021
bat-SL-CoVZC45
HN2021A
HN2021B
bat-SL-CoVZXC21
HN2021G
BetaCoV_Yunnan_Rp_JCC9_2020
YNLF_31C
YNLF_34C
Rf1
BtRf-SX2013
BtRf-BetaCoV_HeB2013
Jiyuan-84
16BO133
BatCoV_B20-50
LN2020A
LN2020G
JTMC15
LN2020C
LN2020F
LN2020H
LN2020E
LN2020B
BtRf-JL2012
Rt22QB8
Rt22QT36
Rt22QT178
Rt22QT124
Rt22QT48
Rt22QT161
Rt22QT53
Ra22QT27
Rt22QT46
SC2018B
BtRs-HuB2013
BtRs-BetaCoV/HB2020-Q329
BtRl-BetaCoV/SC2018
GZ2021C
GZ2021I
Rt17DN420
Rs4247
BtRs-BetaCoV/YN2018A
Rt21LC92
YN2020A
BtRs-BetaCoV/YN2020-Q322
Rt22SL9
RtVN21-192
Rt22SL85
Rt22SL115
Rt22SL67
Rt22SL130
Rt22SL92
Rt22SL58
RtVN21-200
RtVN21-197
RtVN21-201
GD2017Q
GD2017P
GD2017W
RsGD2014B
RsGD2014A
JX2021L
JX2021N
Rp3
Rt22CB395
RaCH025
Rs56
FJ2021A
JX2021G
FJ2021E
FJ2021M
GD2017G
GD2017M
JX2021M
JX2021J
JX2021O
JX2021P
JX2021AC
HB2020D
RsHuB2019A
RsHuB2019B
GD2016B
GD2017J
GD2017I
GD2017K
GD2017L
GD2017O
GD2017H
GD2017N
AH2021A
FJ2021D
Longquan-140
JX2021K
GD2019E
GD2019D
BtRs-GX2013
HKU3-7
HKU3-8
JX2021C
HB2020E
GD2017F
Ra21CB8
GD2019A
GD2019B
HKU3-12
HKU3-13
HKU3-10
HKU3-3
HKU3-4
HKU3-6
HKU3-2
HKU3-5
HKU3-1
HKU3-9
HKU3-11
0.1

**Figure S1. Phylogenetic tree of all sarbecovirus S RBDs, related to Figure 1**

Maximum likelihood tree of sarbecoviruses based on the nucleotide sequence of their S RBDs. Node support is shown above each branch. Scale bar indicates genetic distance (nucleotide substitutions per site). Distinct clades are annotated based on a previous published work[18], consistent with **Figure 1A**.

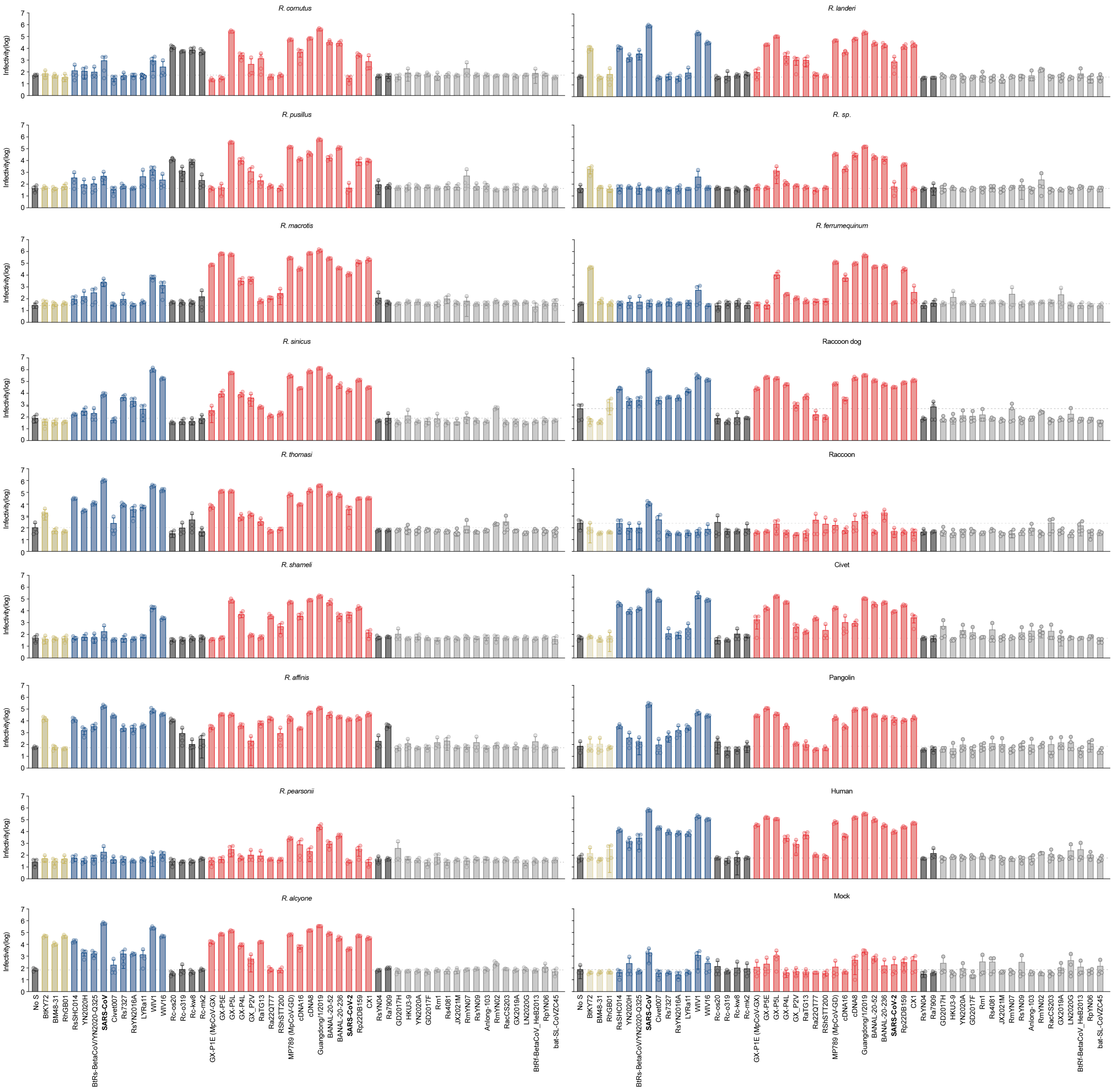


**Figure S2. Round-robin pseudovirus assay of sarbecoviruses and ACE2s, related to Figure 1**

Round-robin pseudovirus assay. HIV-1-based reporter viruses pseudotyped with the S proteins of 53 sarbecoviruses were prepared. The pseudoviruses were inoculated into a series of HOS-TMPRSS2 cells stably expressing *Rhinolophus* bat ACE2 cells at 4 ng HIV-1 p24 antigen. The infectivity (relative light unit) in each target cell is shown. Data are expressed as the mean with SD. Assays were performed in quadruplicate.

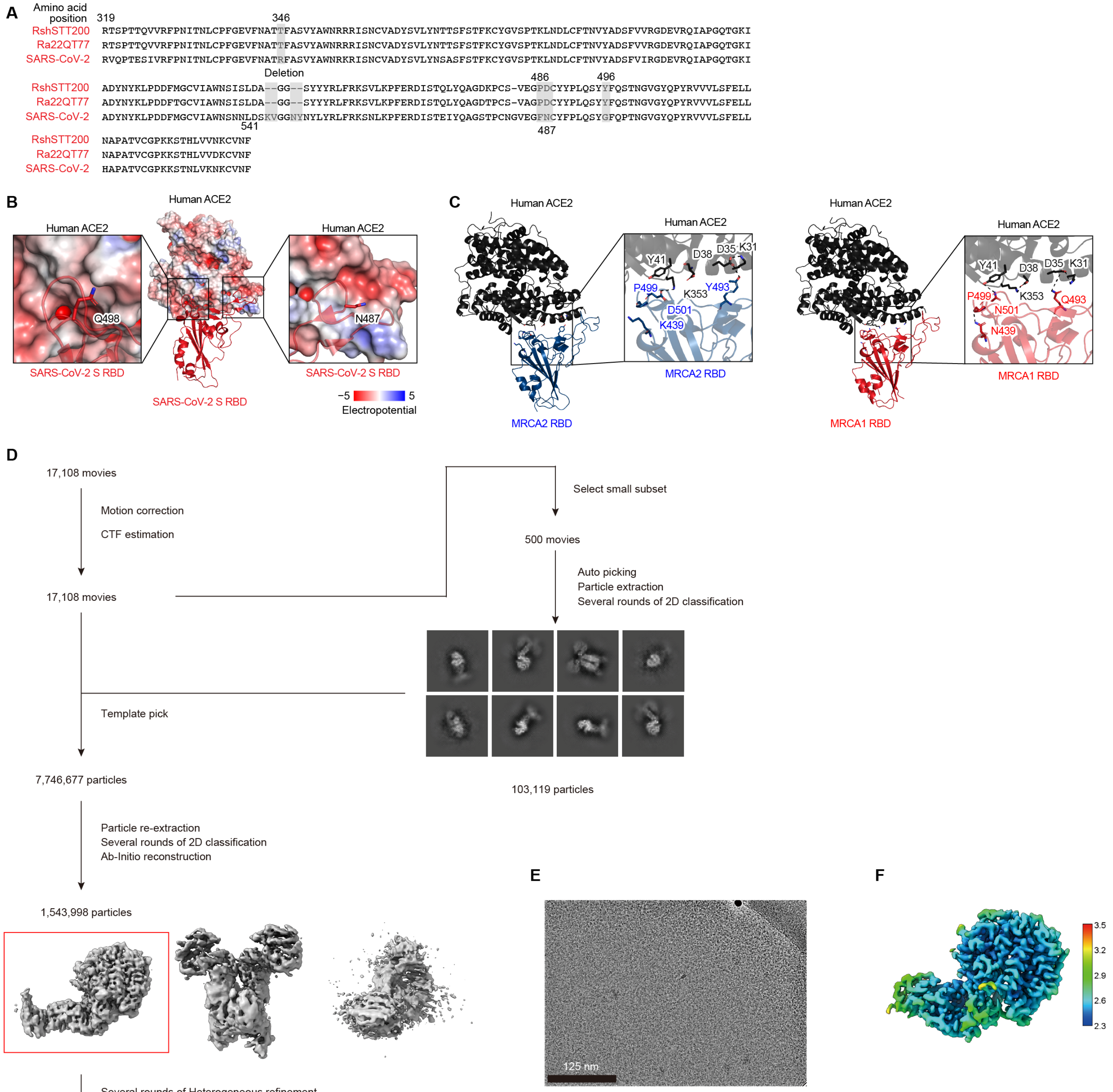


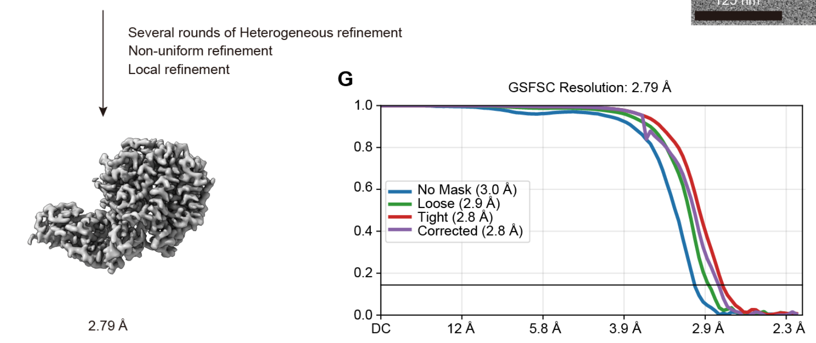


**H**

| | *R. cornutus* ACE2 | Length (Å) | Rc-o319 RBD |
|---|---|---|---|
| Hydrogen bonds | K27 | 2.72 | Q474 |
| | D31 | 2.87 | K487 |
| | | 2.85 | K493 |
| | S34 | 2.87 | Y453 |
| | Y41 | 2.61 | S500(OG) |
| | Q42 | 2.70 | Y498 |
| | E75 | 3.01 | S500 |
| | K353 | 3.41 | H505 |
| | | 2.71 | G502 |
| | | 3.20 | G496 |
| | NAG (N38) | 2.81 | N494 |
| Salt bridges | D31 | 2.67 | K487 |
| | | 2.85 | K493 |
| Hydrophobic bond | F28 | | V475 |
| | Y41 | | Y498 |
| | | | A501 |
| | L45 | | Y498 |

**Figure S3. The impact of amino acid difference for human ACE2 usage and Cryo-EM data of the costructure of Rc-o319 S RBD and *R. cornutus* ACE2, related to Figures 2 and 3**

(**A**) Amino acid alignment of the RBDs of RshSTT200, Ra22QT77 and SARS-CoV-2. Residues with nonsynonymous substitution or deletion focused on in this study are shaded in gray.

(**B**) Negative charged surface of the human ACE2 spatially proximate to residues 487 and 498 of SARS-CoV-2 S (red) (PDB: 6M0J)[30]. The RBD structure is shown as a cartoon, the ACE2 structure is shown as a surface with electropotential, and the residues 487 and 498 in RBD is shown as a stick.

(**C**) Structural insights into the binding of S RBD and ACE2 proteins. (left) The structural model of complex of MRCA2 RBD (blue) and the human ACE2 (black). (right) The structural model of complex of MRCA2 RBD (red) and the human ACE2 (black). These models were estimated by AlphaFold3. Residues 439, 493, 499 and 501 of S RBD and residues 31, 35, 38, 41 and 353 of ACE2s are indicated as stick model. Dashed lines indicate salt bridges or hydrogen bonds.

(**D-G**) Cryo-EM data collection and refinement of the costructure of Rc-o319 S RBD and *R. cornutus* ACE2.

(**D**) Data processing flowchart. CTF: contrast transfer function.

(**E**) Representative electron micrograph of the costructure of Rc-o319 S RBD and *R. cornutus* ACE2 embedded in vitreous ice. The scale bar represents 125 nm.

(**F**) Local resolution map calculated using CryoSPARC and plotted onto the sharpened cryo-EM map.

(**G**) Global resolution assessment of cryo-EM maps by Gold Standard Fourier shell correlation (GSFSC) curves at the 0.143 criteria.

(**H**) The interaction between Rc-o319 S RBD and *R. cornutus* ACE2. The residue numbering is based on SARS-CoV-2.

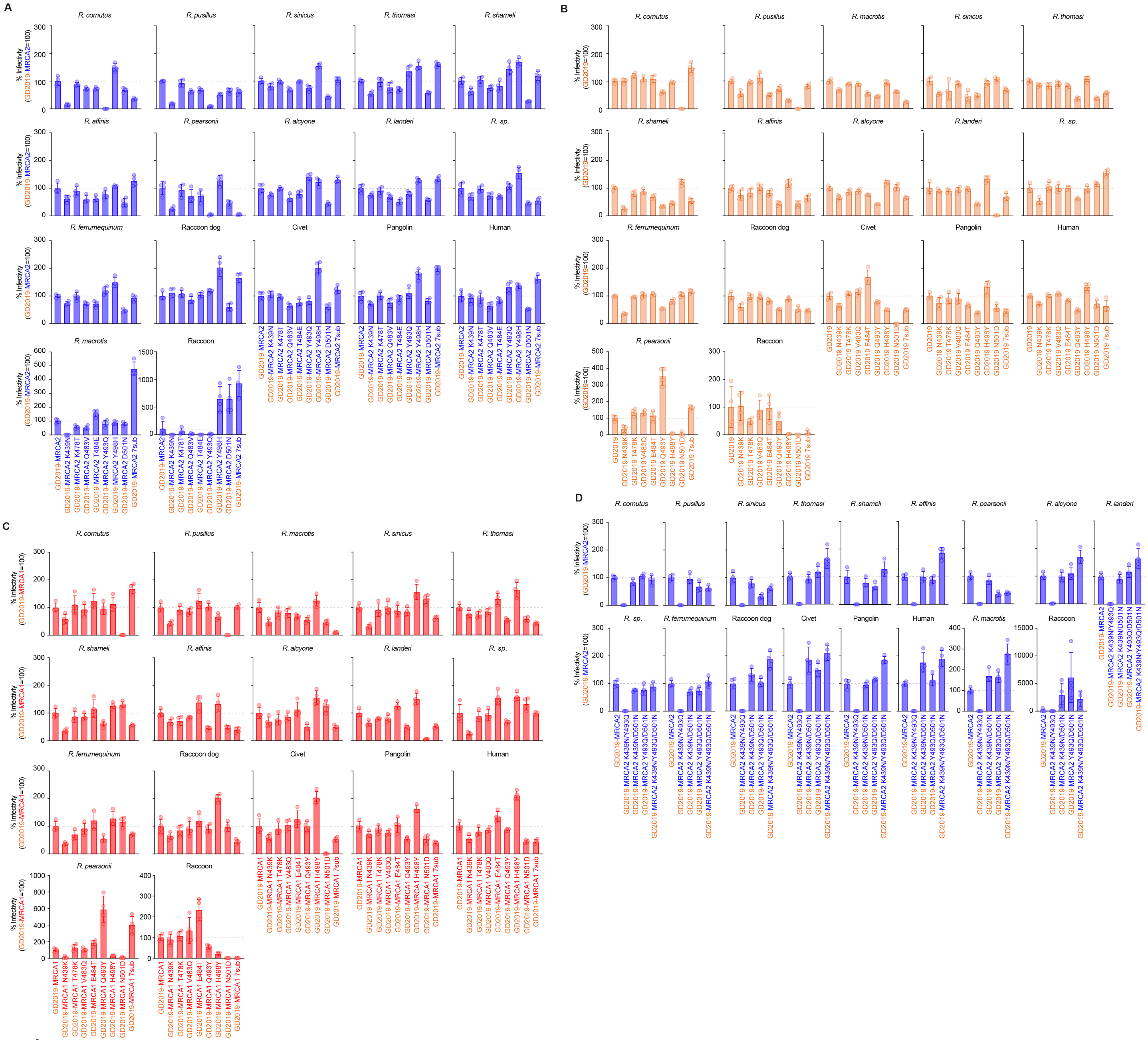


**Figure S4. Pseudovirus assay of chimera S protein, related to Figure 4**

(**A-D**) Pseudovirus assay. HIV-1-based reporter viruses pseudotyped with GD2019-MRCA2 (**A, D**), GD2019 (**B**) or GD2019-MRCA1 (**C**) protein, or the derivatives were prepared. The pseudoviruses were inoculated into a series of HOS-TMPRSS2 cells stably expressing *Rhinolophus* bat ACE2 cells at 4 ng HIV-1 p24 antigen. Heatmap plotted by the mean values (quadruplicates of independently bound or infected cells) which are representative results out of three independent experiments.

**Table S1. Sarbecovirus S sequences used in this study, related to Figure 1**

| Name | Accession number | Host species | Source | Clade |
|---|---|---|---|---|
| BtRs-BetaCoV/YN2018B | MK211376.1 | *Rhinolophus sinicus* | 10.3389/fmicb.2019.01900 | Clade 1a |
| BtRs-BetaCoV/YN2020-Q324 | OQ175344.1 | *Rhinolophus sinicus* | 10.1038/s41467-023-41264-z | Clade 1a |
| **BtRs-BetaCoV/YN2020-Q325** | **OQ175345.1** | ***Rhinolophus sinicus*** | **10.1038/s41467-023-41264-z** | **Clade 1a** |
| BtRs-BetaCoV/YN2020-Q326 | OQ175346.1 | *Rhinolophus sinicus* | 10.1038/s41467-023-41264-z | Clade 1a |
| BtRs-BetaCoV/YN2020-Q327 | OQ175347.1 | *Rhinolophus sinicus* | 10.1038/s41467-023-41264-z | Clade 1a |
| BtRs-BetaCoV/YN2020-Q328 | OQ175348.1 | *Rhinolophus sinicus* | 10.1038/s41467-023-41264-z | Clade 1a |
| **Civet007** | **AY572034.1** | ***Paguma larvata*** | **10.3201/eid1112.041293** | **Clade 1a** |
| LS1 | OP963575.1 | *Rhinolophus thomasi* | 10.1038/s41467-023-39835-1 | Clade 1a |
| **LYRa11** | **KF569996.1** | ***Rhinolophus affinis*** | **10.1128/JVI.00631-14** | **Clade 1a** |
| Rs3367 | KC881006.1 | *Rhinolophus sinicus* | 10.1038/nature12711 | Clade 1a |
| Rs4084 | KY417144.1 | *Rhinolophus sinicus* | 10.1371/journal.ppat.1006698 | Clade 1a |
| Rs4231 | KY417146.1 | *Rhinolophus sinicus* | 10.1371/journal.ppat.1006698 | Clade 1a |
| Rs4874 | KY417150.1 | *Rhinolophus sinicus* | 10.1371/journal.ppat.1006698 | Clade 1a |
| **Rs7327** | **KY417151.1** | ***Rhinolophus sinicus*** | **10.1371/journal.ppat.1006698** | **Clade 1a** |
| Rs9401 | KY417152.1 | *Rhinolophus sinicus* | 10.1371/journal.ppat.1006698 | Clade 1a |
| **RsSHC014** | **KC881005.1** | ***Rhinolophus sinicus*** | **10.1038/nature12711** | **Clade 1a** |
| RsYN2012 | OQ503500.1 | *Rhinolophus sinicus* | 10.1128/jvi.00395-23 | Clade 1a |
| RsYN2013 | OQ503501.1 | *Rhinolophus sinicus* | 10.1128/jvi.00395-23 | Clade 1a |
| **RsYN2016A** | **OQ503504.1** | ***Rhinolophus sinicus*** | **10.1128/jvi.00395-23** | **Clade 1a** |
| RsYN2016C | OQ503505.1 | *Rhinolophus sinicus* | 10.1128/jvi.00395-23 | Clade 1a |
| **SARS-CoV** | **AY394995.1** | ***Homo sapiens*** | **10.1126/science.1092002** | **Clade 1a** |
| **WIV1** | **KF367457.1** | ***Rhinolophus sinicus*** | **10.1038/nature12711** | **Clade 1a** |
| **WIV16** | **KT444582.1** | ***Rhinolophus sinicus*** | **10.1128/JVI.02582-15** | **Clade 1a** |
| YN2016A | OK017847.1 | *Rhinolophus sinicus* | 10.1093/nsr/nwac213 | Clade 1a |
| YN2016B | OK017848.1 | *Rhinolophus sinicus* | 10.1093/nsr/nwac213 | Clade 1a |
| YN2016C | OK017849.1 | *Rhinolophus sinicus* | 10.1093/nsr/nwac213 | Clade 1a |
| YN2016D | OK017850.1 | *Rhinolophus sinicus* | 10.1093/nsr/nwac213 | Clade 1a |
| YN2016E | OK017851.1 | *Rhinolophus sinicus* | 10.1093/nsr/nwac213 | Clade 1a |
| YN2020B | OK017852.1 | *Rhinolophus sinicus* | 10.1093/nsr/nwac213 | Clade 1a |
| YN2020C | OK017853.1 | *Rhinolophus sinicus* | 10.1093/nsr/nwac213 | Clade 1a |
| YN2020D | OK017854.1 | *Rhinolophus sinicus* | 10.1093/nsr/nwac213 | Clade 1a |
| YN2020E | OK017855.1 | *Rhinolophus sinicus* | 10.1093/nsr/nwac213 | Clade 1a |
| YN2020F | OK017856.1 | *Rhinolophus sinicus* | 10.1093/nsr/nwac213 | Clade 1a |
| YN2020G | OK017857.1 | *Rhinolophus sinicus* | 10.1093/nsr/nwac213 | Clade 1a |
| **YN2020H** | **OK017858.1** | ***Rhinolophus sinicus*** | **10.1093/nsr/nwac213** | **Clade 1a** |
| BANAL-20-103 | MZ937001.1 | *Rhinolophus pusillus* | 10.1038/s41586-022-04532-4 | Clade 1b |
| **BANAL-20-236** | **MZ937003.1** | ***Rhinolophus marshalli*** | **10.1038/s41586-022-04532-4** | **Clade 1b** |
| **BANAL-20-52** | **MZ937000.1** | ***Rhinolophus malayanus*** | **10.1038/s41586-022-04532-4** | **Clade 1b** |
| **cDNA16** | **MT799523.1** | ***Manis javanica*** | **10.1038/s41586-020-2313-x** | **Clade 1b** |
| **cDNA8** | **MT799521.1** | ***Manis javanica*** | **10.1038/s41586-020-2313-x** | **Clade 1b** |
| **CX1** | **OP963576.1** | ***Rhinolophus pusillus*** | **10.1038/s41467-023-39835-1** | **Clade 1b** |
| **Guangdong/1/2019** | **EPI_ISL_410721** | ***Manis javanica*** | **10.1038/s41586-020-2313-x** | **Clade 1b** |
| **GX-P1E (MpCoV-GX)** | **MT040334.1** | ***Manis javanica*** | **10.1038/s41586-020-2169-0** | **Clade 1b** |
| **GX-P2V** | **MT072864.1** | ***Manis javanica*** | **10.1038/s41586-020-2169-0** | **Clade 1b** |
| **GX-P4L** | **MT040333.1** | ***Manis javanica*** | **10.1038/s41586-020-2169-0** | **Clade 1b** |
| **GX-P5E** | **MT040336.1** | ***Manis javanica*** | **10.1038/s41586-020-2169-0** | **Clade 1b** |
| **GX-P5L** | **MT040335.1** | ***Manis javanica*** | **10.1038/s41586-020-2169-0** | **Clade 1b** |
| **MP789 (MpCoV-GD)** | **MT121216.1** | ***Manis javanica*** | **10.1371/journal.ppat.1008421** | **Clade 1b** |
| Ra22QT106 | OR233322.1 | *Rhinolophus affinis* | 10.1111/mec.17486 | Clade 1b |
| Ra22QT135 | OR233323.1 | *Rhinolophus affinis* | 10.1111/mec.17486 | Clade 1b |
| Ra22QT137 | OR233328.1 | *Rhinolophus affinis* | 10.1111/mec.17486 | Clade 1b |
| **Ra22QT77** | **OR233324.1** | ***Rhinolophus affinis*** | **10.1111/mec.17486** | **Clade 1b** |
| **RaTG13** | **MN996532.2** | ***Rhinolophus affinis*** | **10.1038/s41586-020-2012-7** | **Clade 1b** |
| **Rp22DB159** | **OR233302.1** | ***Rhinolophus pusillus*** | **10.1111/mec.17486** | **Clade 1b** |
| RshSTT182 | EPI_ISL_852604 | *Rhinolophus shameli* | 10.1101/2021.01.26.428212 | Clade 1b |
| **RshSTT200** | **EPI_ISL_852605** | ***Rhinolophus shameli*** | **10.1101/2021.01.26.428212** | **Clade 1b** |
| **SARS-CoV-2** | **NC_045512.2** | ***Homo sapiens*** | **10.1038/s41586-020-2008-3** | **Clade 1b** |
| 16BO133 | KY938558.1 | *Rhinolophus ferrumequinum* | 10.1007/s00248-017-1033-8 | Clade 2 |
| AH2021A | OK017807.1 | *Rhinolophus sinicus* | 10.1093/nsr/nwac213 | Clade 2 |
| **Anlong-103** | **KY770858.1** | ***Rhinolophus sinicus*** | **10.1016/j.virol.2017.03.019** | **Clade 2** |
| Anlong-112 | KY770859.1 | *Rhinolophus sinicus* | 10.1016/j.virol.2017.03.019 | Clade 2 |
| As6526 | KY417142.1 | *Aselliscus stoliczkanus* | 10.1371/journal.ppat.1006698 | Clade 2 |
| BANAL-20-116 | MZ937002.1 | *Rhinolophus malayanus* | 10.1038/s41586-022-04532-4 | Clade 2 |
| BANAL-20-247 | MZ937004.1 | *Rhinolophus malayanus* | 10.1038/s41586-022-04532-4 | Clade 2 |
| **bat-SL-CoVZC45** | **MG772933.1** | ***Rhinolophus pusillus*** | **10.1038/s41426-018-0155-5** | **Clade 2** |
| bat-SL-CoVZXC21 | MG772934.1 | *Rhinolophus pusillus* | 10.1038/s41426-018-0155-5 | Clade 2 |
| BatCoV_B20-50 | ON378802.1 | *Rhinolophus ferrumequinum* | 10.3390/v14071389 | Clade 2 |
| BetaCoV_Yunnan_Rp_JCC9_2020 | OK287355.1 | *Rhinolophus blythi* | 10.1186/s12917-024-04310-6 | Clade 2 |
| BetaCoV_Yunnan_Rs_KMC6_2020 | OK287354.1 | *Rhinolophus sinicus* | 10.1186/s12917-024-04310-6 | Clade 2 |
| BtCoV/279/2005 | DQ648857.1 | *Rhinolophus macrotis* | 10.1128/JVI.00697-06 | Clade 2 |
| **BtRf-BetaCoV_HeB2013** | **KJ473812.1** | ***Rhinolophus ferrumequinum*** | **10.1093/infdis/jiv476** | **Clade 2** |

| | | | | |
|---|---|---|---|---|
| BtRf-JL2012 | KJ473811.1 | *Rhinolophus ferrumequinum* | 10.1093/infdis/jiv476 | Clade 2 |
| BtRf-SX2013 | KJ473813.1 | *Rhinolophus ferrumequinum* | 10.1093/infdis/jiv476 | Clade 2 |
| BtRl-BetaCoV/SC2018 | MK211374.1 | *Rhinolophus sp.* | 10.3389/fmicb.2019.01900 | Clade 2 |
| BtRp-BetaCoV/GD2018-Q323 | OQ175343.1 | *Rhinolophus pusillus* | 10.1038/s41467-023-41264-z | Clade 2 |
| BtRs-BetaCoV/HB2020-Q329 | OQ175349.1 | *Rhinolophus sinicus* | 10.1038/s41467-023-41264-z | Clade 2 |
| BtRs-BetaCoV/YN2018A | MK211375.1 | *Rhinolophus sinicus* | 10.3389/fmicb.2019.01900 | Clade 2 |
| BtRs-BetaCoV/YN2018C | MK211377.1 | *Rhinolophus sinicus* | 10.3389/fmicb.2019.01900 | Clade 2 |
| BtRs-BetaCoV/YN2018D | MK211378.1 | *Rhinolophus sinicus* | 10.3389/fmicb.2019.01900 | Clade 2 |
| BtRs-BetaCoV/YN2020-Q322 | OQ175342.1 | *Rhinolophus affinis* | 10.1038/s41467-023-41264-z | Clade 2 |
| BtRs-BetaCoV/YN2020-Q330 | OQ175350.1 | *Rhinolophus sinicus* | 10.1038/s41467-023-41264-z | Clade 2 |
| BtRs-GX2013 | KJ473815.1 | *Rhinolophus sinicus* | 10.1093/infdis/jiv476 | Clade 2 |
| BtRs-HuB2013 | KJ473814.1 | *Rhinolophus sinicus* | 10.1038/ismej.2015.138 | Clade 2 |
| BtRs-YN2013 | KJ473816.1 | *Rhinolophus sinicus* | 10.1093/infdis/jiv476 | Clade 2 |
| Cp/Yunnan2011 | JX993988.1 | *Chaerephon plicata* | 10.3201/eid1906.121648 | Clade 2 |
| F46 | KU973692.1 | *Rhinolophus pusillus* | 10.1038/emi.2016.140 | Clade 2 |
| FJ2021A | OK017808.1 | *Rhinolophus sinicus* | 10.1093/nsr/nwac213 | Clade 2 |
| FJ2021D | OK017809.1 | *Rhinolophus sinicus* | 10.1093/nsr/nwac213 | Clade 2 |
| FJ2021E | OK017810.1 | *Rhinolophus sinicus* | 10.1093/nsr/nwac213 | Clade 2 |
| FJ2021M | OK017811.1 | *Rhinolophus sinicus* | 10.1093/nsr/nwac213 | Clade 2 |
| GD2016B | OK017812.1 | *Rhinolophus sinicus* | 10.1093/nsr/nwac213 | Clade 2 |
| **GD2017F** | **OK017792.1** | ***Rhinolophus affinis*** | **10.1093/nsr/nwac213** | **Clade 2** |
| GD2017G | OK017813.1 | *Rhinolophus sinicus* | 10.1093/nsr/nwac213 | Clade 2 |
| **GD2017H** | **OK017814.1** | ***Rhinolophus sinicus*** | **10.1093/nsr/nwac213** | **Clade 2** |
| GD2017I | OK017815.1 | *Rhinolophus sinicus* | 10.1093/nsr/nwac213 | Clade 2 |
| GD2017J | OK017816.1 | *Rhinolophus sinicus* | 10.1093/nsr/nwac213 | Clade 2 |
| GD2017K | OK017817.1 | *Rhinolophus sinicus* | 10.1093/nsr/nwac213 | Clade 2 |
| GD2017L | OK017818.1 | *Rhinolophus sinicus* | 10.1093/nsr/nwac213 | Clade 2 |
| GD2017M | OK017819.1 | *Rhinolophus sinicus* | 10.1093/nsr/nwac213 | Clade 2 |
| GD2017N | OK017820.1 | *Rhinolophus sinicus* | 10.1093/nsr/nwac213 | Clade 2 |
| GD2017O | OK017821.1 | *Rhinolophus sinicus* | 10.1093/nsr/nwac213 | Clade 2 |
| GD2017P | OK017822.1 | *Rhinolophus sinicus* | 10.1093/nsr/nwac213 | Clade 2 |
| GD2017Q | OK017823.1 | *Rhinolophus sinicus* | 10.1093/nsr/nwac213 | Clade 2 |
| GD2017W | OK017824.1 | *Rhinolophus sinicus* | 10.1093/nsr/nwac213 | Clade 2 |
| GD2019A | OK017825.1 | *Rhinolophus sinicus* | 10.1093/nsr/nwac213 | Clade 2 |
| GD2019B | OK017826.1 | *Rhinolophus sinicus* | 10.1093/nsr/nwac213 | Clade 2 |
| GD2019D | OK017827.1 | *Rhinolophus sinicus* | 10.1093/nsr/nwac213 | Clade 2 |
| GD2019E | OK017828.1 | *Rhinolophus sinicus* | 10.1093/nsr/nwac213 | Clade 2 |
| **GX2019A** | **OK017859.1** | ***Rhinolophus siamensis*** | **10.1093/nsr/nwac213** | **Clade 2** |
| GZ2021C | OK017829.1 | *Rhinolophus sinicus* | 10.1093/nsr/nwac213 | Clade 2 |
| GZ2021H | OK017830.1 | *Rhinolophus sinicus* | 10.1093/nsr/nwac213 | Clade 2 |
| GZ2021I | OK017831.1 | *Rhinolophus sinicus* | 10.1093/nsr/nwac213 | Clade 2 |
| HB2020D | OK017801.1 | *Rhinolophus sp.* | 10.1093/nsr/nwac213 | Clade 2 |
| HB2020E | OK017802.1 | *Rhinolophus sp.* | 10.1093/nsr/nwac213 | Clade 2 |
| HKU3-1 | DQ022305.2 | *Rhinolophus sinicus* | 10.1128/JVI.02219-09 | Clade 2 |
| HKU3-10 | GQ153545.1 | *Rhinolophus sinicus* | 10.1128/JVI.02219-09 | Clade 2 |
| HKU3-11 | GQ153546.1 | *Rhinolophus sinicus* | 10.1128/JVI.02219-09 | Clade 2 |
| HKU3-12 | GQ153547.1 | *Rhinolophus sinicus* | 10.1128/JVI.02219-09 | Clade 2 |
| HKU3-13 | GQ153548.1 | *Rhinolophus sinicus* | 10.1128/JVI.02219-09 | Clade 2 |
| HKU3-2 | DQ084199.1 | *Rhinolophus sinicus* | 10.1128/JVI.02219-09 | Clade 2 |
| HKU3-3 | DQ084200.1 | *Rhinolophus sinicus* | 10.1128/JVI.02219-09 | Clade 2 |
| HKU3-4 | GQ153539.1 | *Rhinolophus sinicus* | 10.1128/JVI.02219-09 | Clade 2 |
| HKU3-5 | GQ153540.1 | *Rhinolophus sinicus* | 10.1128/JVI.02219-09 | Clade 2 |
| HKU3-6 | GQ153541.1 | *Rhinolophus sinicus* | 10.1128/JVI.02219-09 | Clade 2 |
| HKU3-7 | GQ153542.1 | *Rhinolophus sinicus* | 10.1128/JVI.02219-09 | Clade 2 |
| HKU3-8 | GQ153543.1 | *Rhinolophus sinicus* | 10.1128/JVI.02219-09 | Clade 2 |
| **HKU3-9** | **GQ153544.1** | ***Rhinolophus sinicus*** | **10.1128/JVI.02219-09** | **Clade 2** |
| HN2021A | OK017803.1 | *Rhinolophus pusillus* | 10.1093/nsr/nwac213 | Clade 2 |
| HN2021B | OK017804.1 | *Rhinolophus pusillus* | 10.1093/nsr/nwac213 | Clade 2 |
| HN2021C | OK017832.1 | *Rhinolophus sinicus* | 10.1093/nsr/nwac213 | Clade 2 |
| HN2021D | OK017833.1 | *Rhinolophus sinicus* | 10.1093/nsr/nwac213 | Clade 2 |
| HN2021E | OK017834.1 | *Rhinolophus sinicus* | 10.1093/nsr/nwac213 | Clade 2 |
| HN2021F | OK017835.1 | *Rhinolophus sinicus* | 10.1093/nsr/nwac213 | Clade 2 |
| HN2021G | OK017805.1 | *Rhinolophus pusillus* | 10.1093/nsr/nwac213 | Clade 2 |
| Jiyuan-84 | KY770860.1 | *Rhinolophus ferrumequinum* | 10.1016/j.virol.2017.03.019 | Clade 2 |
| JTMC15 | KU182964.1 | *Rhinolophus ferrumequinum* | 10.1007/s12250-016-3727-3 | Clade 2 |
| JX2021AC | OK017836.1 | *Rhinolophus sinicus* | 10.1093/nsr/nwac213 | Clade 2 |
| JX2021C | OK017837.1 | *Rhinolophus sinicus* | 10.1093/nsr/nwac213 | Clade 2 |
| JX2021D | OK017860.1 | *Rhinolophus siamensis* | 10.1093/nsr/nwac213 | Clade 2 |
| JX2021G | OK017838.1 | *Rhinolophus sinicus* | 10.1093/nsr/nwac213 | Clade 2 |
| JX2021J | OK017839.1 | *Rhinolophus sinicus* | 10.1093/nsr/nwac213 | Clade 2 |
| JX2021K | OK017840.1 | *Rhinolophus sinicus* | 10.1093/nsr/nwac213 | Clade 2 |
| JX2021L | OK017841.1 | *Rhinolophus sinicus* | 10.1093/nsr/nwac213 | Clade 2 |
| **JX2021M** | **OK017842.1** | ***Rhinolophus sinicus*** | **10.1093/nsr/nwac213** | **Clade 2** |

| | | | | |
|---|---|---|---|---|
| JX2021N | OK017843.1 | *Rhinolophus sinicus* | 10.1093/nsr/nwac213 | Clade 2 |
| JX2021O | OK017844.1 | *Rhinolophus sinicus* | 10.1093/nsr/nwac213 | Clade 2 |
| JX2021P | OK017845.1 | *Rhinolophus sinicus* | 10.1093/nsr/nwac213 | Clade 2 |
| LN2020A | OK017794.1 | *Rhinolophus ferrumequinum* | 10.1093/nsr/nwac213 | Clade 2 |
| LN2020B | OK017795.1 | *Rhinolophus ferrumequinum* | 10.1093/nsr/nwac213 | Clade 2 |
| LN2020C | OK017796.1 | *Rhinolophus ferrumequinum* | 10.1093/nsr/nwac213 | Clade 2 |
| LN2020E | OK017797.1 | *Rhinolophus ferrumequinum* | 10.1093/nsr/nwac213 | Clade 2 |
| LN2020F | OK017798.1 | *Rhinolophus ferrumequinum* | 10.1093/nsr/nwac213 | Clade 2 |
| **LN2020G** | **OK017799.1** | ***Rhinolophus ferrumequinum*** | **10.1093/nsr/nwac213** | **Clade 2** |
| LN2020H | OK017800.1 | *Rhinolophus ferrumequinum* | 10.1093/nsr/nwac213 | Clade 2 |
| Longquan-140 | KF294457.1 | *Rhinolophus monoceros* | 10.1016/j.virol.2017.03.019 | Clade 2 |
| PrC31 | MW703458.1 | *Rhinolophus pusillus* | 10.1080/22221751.2021.1964925 | Clade 2 |
| Ra21CB8 | OR233291.1 | *Rhinolophus affinis* | 10.1111/mec.17486 | Clade 2 |
| Ra22DB107 | OR233292.1 | *Rhinolophus affinis* | 10.1111/mec.17486 | Clade 2 |
| Ra22DB191 | OR233304.1 | *Rhinolophus affinis* | 10.1111/mec.17486 | Clade 2 |
| Ra22QT27 | OR233315.1 | *Rhinolophus affinis* | 10.1111/mec.17486 | Clade 2 |
| **RacCS203** | **MW251308.1** | ***Rhinolophus acuminatus*** | **10.1038/s41467-021-21240-1** | **Clade 2** |
| RaCH025 | OM240725.1 | *Rhinolophus affinis* | 10.1128/mbio.00463-22 | Clade 2 |
| Rf1 | DQ412042.1 | *Rhinolophus ferrumequinum* | 10.1126/science.1118391 | Clade 2 |
| Rf4092 | KY417145.1 | *Rhinolophus ferrumequinum* | 10.1371/journal.ppat.1006698 | Clade 2 |
| **Rm1** | **DQ412043.1** | ***Rhinolophus macrotis*** | **10.1126/science.1118391** | **Clade 2** |
| **RmYN02** | **EPI_ISL_412977** | ***Rhinolophus malayanus*** | **10.1016/j.cub.2020.05.023** | **Clade 2** |
| **RmYN07** | **MZ081377.1** | ***Rhinolophus malayanus*** | **10.1016/j.cell.2021.06.008** | **Clade 2** |
| Rp/Shaanxi2011 | JX993987.1 | *Rhinolophus pusillus* | 10.3201/eid1906.121648 | Clade 2 |
| Rp3 | DQ071615.1 | *Rhinolophus pearsoni* | 10.1126/science.1118391 | Clade 2 |
| **RpYN06** | **MZ081381.1** | ***Rhinolophus pusillus*** | **10.1016/j.cell.2021.06.008** | **Clade 2** |
| **Rs4081** | **KY417143.1** | ***Rhinolophus sinicus*** | **10.1038/nature12711** | **Clade 2** |
| Rs4237 | KY417147.1 | *Rhinolophus sinicus* | 10.1371/journal.ppat.1006698 | Clade 2 |
| Rs4247 | KY417148.1 | *Rhinolophus sinicus* | 10.1371/journal.ppat.1006698 | Clade 2 |
| Rs4255 | KY417149.1 | *Rhinolophus sinicus* | 10.1371/journal.ppat.1006698 | Clade 2 |
| Rs56 | MW681002.1 | *Rhinolophus sinicus* | 10.1128/mbio.00463-22 | Clade 2 |
| Rs672 | FJ588686.1 | *Rhinolophus sinicus* | 10.1099/vir.0.016378-0 | Clade 2 |
| RsGD2014A | OQ503495.1 | *Rhinolophus sinicus* | 10.1128/jvi.00395-23 | Clade 2 |
| RsGD2014B | OQ503496.1 | *Rhinolophus sinicus* | 10.1128/jvi.00395-23 | Clade 2 |
| RsGZ2015 | OQ503497.1 | *Rhinolophus sinicus* | 10.1128/jvi.00395-23 | Clade 2 |
| RsHuB2019A | OQ503498.1 | *Rhinolophus sinicus* | 10.1128/jvi.00395-23 | Clade 2 |
| RsHuB2019B | OQ503499.1 | *Rhinolophus sinicus* | 10.1128/jvi.00395-23 | Clade 2 |
| RsVN21-195 | OR261265.1 | *Rhinolophus siamensis* | 10.3390/v15091897 | Clade 2 |
| RsYN03 | MZ081379.1 | *Rhinolophus sinicus* | 10.1016/j.cell.2021.06.008 | Clade 2 |
| **RsYN09** | **MZ081382.1** | ***Rhinolophus stheno*** | **10.1016/j.cell.2021.06.008** | **Clade 2** |
| RsYN2014 | OQ503502.1 | *Rhinolophus sinicus* | 10.1128/jvi.00395-23 | Clade 2 |
| RsYN2018 | OQ503506.1 | *Rhinolophus thomasi* | 10.1128/jvi.00395-23 | Clade 2 |
| Rt17DN420 | OR233295.1 | *Rhinolophus thomasi* | 10.1111/mec.17486 | Clade 2 |
| Rt21LC39 | OR233308.1 | *Rhinolophus thomasi* | 10.1111/mec.17486 | Clade 2 |
| Rt21LC92 | OR233297.1 | *Rhinolophus thomasi* | 10.1111/mec.17486 | Clade 2 |
| Rt22CB395 | OR233293.1 | *Rhinolophus thomasi* | 10.1111/mec.17486 | Clade 2 |
| Rt22DB31 | OR233294.1 | *Rhinolophus thomasi* | 10.1111/mec.17486 | Clade 2 |
| Rt22DB38 | OR233310.1 | *Rhinolophus thomasi* | 10.1111/mec.17486 | Clade 2 |
| Rt22LC371 | OR233306.1 | *Rhinolophus thomasi* | 10.1111/mec.17486 | Clade 2 |
| Rt22LC376 | OR233307.1 | *Rhinolophus thomasi* | 10.1111/mec.17486 | Clade 2 |
| Rt22LC378 | OR233296.1 | *Rhinolophus thomasi* | 10.1111/mec.17486 | Clade 2 |
| Rt22QB78 | OR233298.1 | *Rhinolophus thomasi* | 10.1111/mec.17486 | Clade 2 |
| Rt22QB8 | OR233299.1 | *Rhinolophus thomasi* | 10.1111/mec.17486 | Clade 2 |
| Rt22QT124 | OR233300.1 | *Rhinolophus thomasi* | 10.1111/mec.17486 | Clade 2 |
| Rt22QT161 | OR233316.1 | *Rhinolophus thomasi* | 10.1111/mec.17486 | Clade 2 |
| Rt22QT178 | OR233317.1 | *Rhinolophus thomasi* | 10.1111/mec.17486 | Clade 2 |
| Rt22QT36 | OR233318.1 | *Rhinolophus thomasi* | 10.1111/mec.17486 | Clade 2 |
| Rt22QT46 | OR233319.1 | *Rhinolophus thomasi* | 10.1111/mec.17486 | Clade 2 |
| Rt22QT48 | OR233320.1 | *Rhinolophus thomasi* | 10.1111/mec.17486 | Clade 2 |
| Rt22QT53 | OR233321.1 | *Rhinolophus thomasi* | 10.1111/mec.17486 | Clade 2 |
| Rt22SL115 | OR233301.1 | *Rhinolophus thomasi* | 10.1111/mec.17486 | Clade 2 |
| Rt22SL130 | OR233311.1 | *Rhinolophus thomasi* | 10.1111/mec.17486 | Clade 2 |
| Rt22SL58 | OR233312.1 | *Rhinolophus thomasi* | 10.1111/mec.17486 | Clade 2 |
| Rt22SL67 | OR233309.1 | *Rhinolophus thomasi* | 10.1111/mec.17486 | Clade 2 |
| Rt22SL85 | OR233313.1 | *Rhinolophus thomasi* | 10.1111/mec.17486 | Clade 2 |
| Rt22SL9 | OR233303.1 | *Rhinolophus thomasi* | 10.1111/mec.17486 | Clade 2 |
| Rt22SL92 | OR233314.1 | *Rhinolophus thomasi* | 10.1111/mec.17486 | Clade 2 |
| Rt321LC192 | OR233305.1 | *Rhinolophus thomasi* | 10.1111/mec.17486 | Clade 2 |
| RtVN21-192 | OR261263.1 | *Rhinolophus thomasi* | 10.3390/v15091897 | Clade 2 |
| RtVN21-193 | OR261264.1 | *Rhinolophus thomasi* | 10.3390/v15091897 | Clade 2 |
| RtVN21-197 | OR261266.1 | *Rhinolophus thomasi* | 10.3390/v15091897 | Clade 2 |
| RtVN21-200 | OR261267.1 | *Rhinolophus thomasi* | 10.3390/v15091897 | Clade 2 |
| RtVN21-201 | OR261268.1 | *Rhinolophus thomasi* | 10.3390/v15091897 | Clade 2 |

| | | | | |
|---|---|---|---|---|
| RtVN21-29 | OR261262.1 | *Rhinolophus thomasi* | 10.3390/v15091897 | Clade 2 |
| SC2018B | OK017846.1 | *Rhinolophus sinicus* | 10.1093/nsr/nwac213 | Clade 2 |
| **YN2020A** | **OK017793.1** | ***Rhinolophus affinis*** | **10.1093/nsr/nwac213** | **Clade 2** |
| YN2021 | OK017806.1 | *Rhinolophus pusillus* | 10.1093/nsr/nwac213 | Clade 2 |
| YNLF_31C | KP886808.1 | *Rhinolophus ferrumequinum* | 10.1128/jvi.01048-15 | Clade 2 |
| YNLF_34C | KP886809.1 | *Rhinolophus ferrumequinum* | 10.1128/jvi.01048-15 | Clade 2 |
| **BM48-31** | **NC_014470.1** | ***Rhinolophus blasii*** | **10.1128/JVI.00650-10** | **Clade 3** |
| **BtKY72** | **KY352407.1** | ***Rhinolophus sp.*** | **10.1128/MRA.00548-19** | **Clade 3** |
| Khosta-1 | MZ190137.1 | *Rhinolophus ferrumequinum* | 10.3390/v14010113 | Clade 3 |
| Khosta-2 | MZ190138.1 | *Rhinolophus hipposideros* | 10.3390/v14010113 | Clade 3 |
| PDF-2370 | MT726044.1 | *Rhinolophus sp.* | 10.1093/ve/veab007 | Clade 3 |
| PDF-2386 | MT726043.1 | *Rhinolophus sp.* | 10.1093/ve/veab007 | Clade 3 |
| PRD-0038 | MT726045.1 | *Rhinolophus sp.* | 10.1093/ve/veab007 | Clade 3 |
| RfGB01 | OQ401247.1 | *Rhinolophus ferrumequinum* | 10.1038/s41467-023-38717-w | Clade 3 |
| RfGB02 | OQ401249.1 | *Rhinolophus ferrumequinum* | 10.1038/s41467-023-38717-w | Clade 3 |
| **RhGB01** | **MW719567.1** | ***Rhinolophus hipposideros*** | **10.1038/s41598-021-94011-z** | **Clade 3** |
| RhGB02 | OP776338.1 | *Rhinolophus hipposideros* | 10.1101/2023.02.14.528476 | Clade 3 |
| RhGB05 | OP776339.1 | *Rhinolophus hipposideros* | 10.1101/2023.02.14.528476 | Clade 3 |
| RhGB06 | OP776340.1 | *Rhinolophus hipposideros* | 10.1101/2023.02.14.528476 | Clade 3 |
| RhGB07 | OQ401248.1 | *Rhinolophus hipposideros* | 10.1038/s41467-023-38717-w | Clade 3 |
| **Rc-kw8** | **LC663793.1** | ***Rhinolophus cornutus*** | **10.3201/eid2812.220803** | **Rc clade** |
| **Rc-mk2** | **LC663959.1** | ***Rhinolophus cornutus*** | **10.3201/eid2812.220801** | **Rc clade** |
| **Rc-o319** | **LC556375.1** | ***Rhinolophus cornutus*** | **10.3201/eid2612.203386** | **Rc clade** |
| **Rc-os20** | **LC663958.1** | ***Rhinolophus cornutus*** | **10.3201/eid2812.220802** | **Rc clade** |
| **Ra7909** | **OL674077.1** | ***Rhinolophus affinis*** | **10.1080/22221751.2021.1956373** | **Yunnan clade** |
| RmYN05 | MZ081376.1 | *Rhinolophus malayanus* | 10.1016/j.cell.2021.06.008 | Yunnan clade |
| RmYN08 | MZ081378.1 | *Rhinolophus malayanus* | 10.1016/j.cell.2021.06.008 | Yunnan clade |
| Rs7896 | GWHBAUM01000001 | *Rhinolophus stheno* | 10.1080/22221751.2021.1956373 | Yunnan clade |
| Rs7905 | GWHBAUN01000001 | *Rhinolophus stheno* | 10.1080/22221751.2021.1956373 | Yunnan clade |
| Rs7907 | GWHBAUO01000001 | *Rhinolophus stheno* | 10.1080/22221751.2021.1956373 | Yunnan clade |
| Rs7921 | GWHBAUQ01000001 | *Rhinolophus stheno* | 10.1080/22221751.2021.1956373 | Yunnan clade |
| Rs7924 | GWHBAUR01000001 | *Rhinolophus stheno* | 10.1080/22221751.2021.1956373 | Yunnan clade |
| Rs7931 | GWHBAUS01000001 | *Rhinolophus stheno* | 10.1080/22221751.2021.1956373 | Yunnan clade |
| Rs7952 | GWHBAUT01000001 | *Rhinolophus stheno* | 10.1080/22221751.2021.1956373 | Yunnan clade |
| **RsYN04** | **MZ081380.1** | ***Rhinolophus stheno*** | **10.1016/j.cell.2021.06.008** | **Yunnan clade** |
| Ra22DB163 | OR233325.1 | *Rhinolophus affinis* | 10.1111/mec.17486 | ND |
| Ra22DB173 | OR233326.1 | *Rhinolophus affinis* | 10.1111/mec.17486 | ND |

Sarbecoviruses used for experiment in this study are indicated as bold.

ND: Not determined

**Table S2. ACE2 sequences used in this study, related to Figure 1**

| Host species | Accession number | Source |
|---|---|---|
| *Rhinolophus affinis* | MT394225.1 | https://www.ncbi.nlm.nih.gov/nuccore/MT394225.1/ |
| *Rhinolophus alcyone* | KR559016.1 | https://www.ncbi.nlm.nih.gov/nuccore/KR559016 |
| *Rhinolophus cornutus* | LC564973 | https://www.ncbi.nlm.nih.gov/nuccore/LC564973 |
| *Rhinolophus ferrumequinum* | AB297479.1 | https://www.ncbi.nlm.nih.gov/nuccore/AB297479.1 |
| *Rhinolophus landeri* | KR559015.1 | https://www.ncbi.nlm.nih.gov/nuccore/KR559015.1 |
| *Rhinolophus macrotis* | GQ999932.1 | https://www.ncbi.nlm.nih.gov/nuccore/GQ999932.1 |
| *Rhinolophus pearsonii* | EF569964.1 | https://www.ncbi.nlm.nih.gov/nuccore/EF569964.1 |
| *Rhinolophus pusillus* | GQ999938.1 | https://www.ncbi.nlm.nih.gov/nuccore/GQ999938.1 |
| *Rhinolophus shameli* | MZ851782.1 | https://www.ncbi.nlm.nih.gov/nuccore/MZ851782.1 |
| *Rhinolophus sinicus* | KC881004.1 | https://www.ncbi.nlm.nih.gov/nuccore/KC881004.1 |
| *Rhinolophus sp.* (PREDICT/PDF-2370) | MW183243.1 | https://www.ncbi.nlm.nih.gov/nuccore/MW183243.1 |
| *Rhinolophus thomasi* | OQ511290.1 | https://www.ncbi.nlm.nih.gov/nuccore/OQ511290.1 |
| Civet | AY881174.1 | https://www.ncbi.nlm.nih.gov/nuccore/AY881174.1 |
| Human | NM_021804.3 | https://www.ncbi.nlm.nih.gov/nuccore/NM_021804.3 |
| Pangolin | XM_017650263.2 | https://www.ncbi.nlm.nih.gov/nuccore/XM_017650263.2 |
| Raccoon | AB211998.1 | https://www.ncbi.nlm.nih.gov/nucleotide/AB211998.1 |
| Raccoon dog | EU024940.1 | https://www.ncbi.nlm.nih.gov/nucleotide/EU024940.1 |

**Table S3. Primers used in this study, related to Figures 2-4**

| Primer name | Primer sequence (5'-to-3') | Purpose |
|---|---|---|
| pC-S BtKY72_Fw | ctatagggcgaattgGGTACCATGAAGTTCTTCATC | Preparation of S expression plasmid |
| pC-S BtKY72_Rv | gagctccaccgcggtgGCGGCCGCTCATGTGTAGTGCAG | Preparation of S expression plasmid |
| BtKY72_K482Q_Fw | GTTACGAACCACTGCAAAGTTACGGGTTCACC | Preparation of S expression plasmid |
| BtKY72_K482Q_Rv | GGTGAACCCGTAACTTTGCAGTGGTTCGTAAC | Preparation of S expression plasmid |
| BtKY72_T487Q_Fw | GAAGAGTTACGGGTTCCAACCAACGGTGGGTGTG | Preparation of S expression plasmid |
| BtKY72_T487Q_Rv | CACACCCACCGTTGGTTGGAACCCGTAACTCTTC | Preparation of S expression plasmid |
| BtKY72_K482Q_T487Q_Fw | GTTACGAACCACTGCAAAGTTACGGGTTCCAACCAACG GTGGGTGTG | Preparation of S expression plasmid |
| BtKY72_K482Q_T487Q_Rv | CACACCCACCGTTGGTTGGAACCCGTAACTTTGCAGTG GTTCGTAAC | Preparation of S expression plasmid |
| Omicron universal Fw | cactatagggcgaattgggtaccatgtttgtgttcctggt | Preparation of S expression plasmid |
| BA.2 WT Rv | agctccaccgcggtggcggccgctcaggtgtagtgcagtttca | Preparation of S expression plasmid |
| CoV-2_Q493K_Fw | CTGTTACTTTCCACTCAAGTCCTATGGCTTCCAAC | Preparation of S expression plasmid |
| CoV-2_Q493K_Rv | GTTGGAAGCCATAGGACTTGAGTGGAAAGTAACAG | Preparation of S expression plasmid |
| CoV-2_Q498T_Fw | CAATCCTATGGCTTCACCCCAACCAATGGAGTG | Preparation of S expression plasmid |
| CoV-2_Q498T_Rv | CACTCCATTGGTTGGGGTGAAGCCATAGGATTG | Preparation of S expression plasmid |
| CoV-2_Q493K_Q498T_Fw | CTGTTACTTTCCACTCAAGTCCTATGGCTTCACCCCAAC CAATGGAGTG | Preparation of S expression plasmid |
| CoV-2_Q493K_Q498T_Rv | CACTCCATTGGTTGGGGTGAAGCCATAGGACTTGAGTGG AAAGTAACAG | Preparation of S expression plasmid |
| pWPI-RcRfACE2-F | ctagcctcgaggtttGGATCCgccaccATGTCAGGCTCTT | Preparation of S expression plasmid |
| pWPI-RcRfACE2-R | agtttaaacACTAGTacgcgtCTACTTGTCATCGTCATCC | Preparation of S expression plasmid |
| RfACE2_D31K_Fw | CAAGAAATTTTTGGACaagTTTAACTCTGAAGCC | Preparation of S expression plasmid |
| RfACE2_D31K_Rv | GGCTTCAGAGTTAAActtGTCCAAAAATTTCTTG | Preparation of S expression plasmid |
| RfACE2_H41Y_Fw | GAAAACCTGTCTtAcCAAAGTTCACTTGCTTC | Preparation of S expression plasmid |
| RfACE2_H41Y_Rv | GAAGCAAGTGAACTTTGgTaAGACAGGTTTTC | Preparation of S expression plasmid |
| pWPI-hACE2-3xFlag-Zeo_Fwd | ctagcctcgaggtttGGATCCgccaccATGTCAagctcttcc | Preparation of S expression plasmid |
| pWPI-3xFlag-Zeo_Common Rev | agtttaaacACTAGTacgcgtttaCTTGTCATCGTC | Preparation of S expression plasmid |
| huACE2_K31D_Fw | caagacatttttggacGACtttaaccacgaagc | Preparation of S expression plasmid |
| huACE2_K31D_Rv | gcttcgtggttaaaGTCgtccaaaaatgtcttg | Preparation of S expression plasmid |
| huACE2_Y41H_Fw | cgaagacctgttcCACcaaagttcacttgcttc | Preparation of S expression plasmid |
| huACE2_Y41H_Rv | gaagcaagtgaactttgGTGgaacaggtcttcg | Preparation of S expression plasmid |
| RShSTT200S Fw | ctatagggcgaattgGGTACCATGATCCTCCTGGCGT | Preparation of S expression plasmid |
| RShSTT200S Rv | agctccaccgcggtgGCGGCCGCTCAGGTATAATGGAG | Preparation of S expression plasmid |
| STT200_T346R_Fw | GTGTTCAACGCCACTAGGTTTGCAAGCGTCTATG | Preparation of S expression plasmid |
| STT200_T346R_Rv | CATAGACGCTTGCAAACCTAGTGGCGTTGAACAC | Preparation of S expression plasmid |
| STT200_P486F_Fw | GCTCCGTGGAGGGTTTCGATTGCTACTATCC | Preparation of S expression plasmid |
| STT200_P486F_Rv | GGATAGTAGCAATCGAAACCCTCCACGGAGC | Preparation of S expression plasmid |
| STT200_D487N_Fw | CCGTGGAGGGTCCGAACTGCTACTATCCGCTTC | Preparation of S expression plasmid |
| STT200_D487N_Rv | GAAGCGGATAGTAGCAGTTCGGACCCTCCACGG | Preparation of S expression plasmid |
| STT200_Y496G_Fw | CCGCTTCAGTCTTACGGCTTTCAATCTACGAAC | Preparation of S expression plasmid |
| STT200_Y498G_Rv | GTTCGTAGATTGAAAGCCGTAAGACTGAAGCGG | Preparation of S expression plasmid |
| STT200_P486F_D487N_Y496G_Fw | GCTCCGTGGAGGGTTTCAACTGCTACTATCCGCTTCAGT CTTACGGC | Preparation of S expression plasmid |
| STT200_P486F_D487N_Y496G_Rv | GTTCGTAGATTGAAAGCCGTAAGACTGAAGCGGATAGTA GCAGTTGAA | Preparation of S expression plasmid |
| CoV-2_del_Fw | CAACAACCTGGACAGCGGCGGTTCTTACCTCTACAGAC TG | Preparation of S expression plasmid |
| CoV-2_del_Rv | CAGTCTGTAGAGGTAAGAACCGCCGCTGTCCAGGTTGTT G | Preparation of S expression plasmid |

| | | |
|---|---|---|
| CoV-2_3mut_Fw | GTAATGGAGTGGAGGGCCCGGATTGTTACTTTCCACTCC AATCCTATTAC | Preparation of S expression plasmid |
| CoV-2_3mut_Rv | CATTGGTTGGTTGGAAGTAATAGGATTGGAGTGGAAAGTA ACAATCCGG | Preparation of S expression plasmid |
| CoV-2_R346T_Fw | GTGTTCAATGCCACCACGTTTGCCTCTGTCTATG | Preparation of S expression plasmid |
| CoV-2_R346T_Rv | CATAGACAGAGGCAAACGTGGTGGCATTGAACAC | Preparation of S expression plasmid |
| CoV-2_F486P_Fw | GTAATGGAGTGGAGGGCCCGAACTGTTACTTTCCAC | Preparation of S expression plasmid |
| CoV-2_F486P_Rv | GTGGAAAGTAACAGTTCGGGCCCTCCACTCCATTAC | Preparation of S expression plasmid |
| CoV-2_N487D_Fw | GGAGTGGAGGGCTTCGATTGTTACTTTCCACTC | Preparation of S expression plasmid |
| CoV-2_N487D_Rv | GAGTGGAAAGTAACAATCGAAGCCCTCCACTCC | Preparation of S expression plasmid |
| CoV-2_G496Y_Fw | CCACTCCAATCCTATTACTTCCAACCAACCAATG | Preparation of S expression plasmid |
| CoV-2_G496Y_Rv | CATTGGTTGGTTGGAAGTAATAGGATTGGAGTGG | Preparation of S expression plasmid |
| QT77_inf_Fw | ctatagggcgaattgGGTACCATGATACTACTAGCG | Preparation of S expression plasmid |
| QT77_inf_Rv | gagctccaccgcggtgGCGGCCGCTCAGGTGTAGTGCAG | Preparation of S expression plasmid |
| QT77_ins_Fw | CATCAGTCTGGATGCAAAGGTGGGAGGTAACTACAACTA CTACTATAGG | Preparation of S expression plasmid |
| QT77_ins_Rv | CCTATAGTAGTAGTTGTAGTTACCTCCCACCTTTGCATCC AGACTGATG | Preparation of S expression plasmid |
| QT77_T346R_Fw | GTTTTCAACGCAACGAgATTCGCATCTGTTTATG | Preparation of S expression plasmid |
| QT77_T346R_Rv | CATAAACAGATGCGAATcTCGTTGCGTTGAAAAC | Preparation of S expression plasmid |
| QT77_P486F_Fw | GTGTAGTGTAGCCGGAttcGATTGTTATTATCCCCTAC | Preparation of S expression plasmid |
| QT77_P486F_Rv | GTAGGGGATAATAACAATCgaaTCCGGCTACACTACAC | Preparation of S expression plasmid |
| QT77_D487N_Fw | GTGTAGTGTAGCCGGACCAaacTGTTATTATCCCCTAC | Preparation of S expression plasmid |
| QT77_D487N_Rv | GTAGGGGATAATAACAgttTGGTCCGGCTACACTACAC | Preparation of S expression plasmid |
| QT77_P486F_D487N_Fw | GTGTAGTGTAGCCGGAttcaacTGTTATTATCCCCTAC | Preparation of S expression plasmid |
| QT77_P486F_D487N_Rv | GTAGGGGATAATAACAgttgaaTCCGGCTACACTACAC | Preparation of S expression plasmid |
| QT77_Y496G_Fw | CCCCTACAGTCTTATggcTTCCAGAGTACCAAC | Preparation of S expression plasmid |
| QT77_Y496G_Rv | GTTGGTACTCTGGAAgccATAAGACTGTAGGGG | Preparation of S expression plasmid |
| o319_infusion_Fw | ctatagggcgaattgGGTACCATGTTTATCTTGGTA | Preparation of S expression plasmid |
| o319_infusion_Rv | gctccaccgcggtgGCGGCCGCTCAGGTATAGTGCAA | Preparation of S expression plasmid |
| o319_Y498A_Fw | CAAGAATTACGGGTTCgcCTCTTCCGCCGGGGAC | Preparation of S expression plasmid |
| o319_Y498A_Rv | GTCCCCGGCGGAAGAGgcGAACCCGTAATTCTTG | Preparation of S expression plasmid |
| o319_S500A_Fw | CAAGAATTACGGGTTCTACTCTgCCGCCGGGGACAGCC ATC | Preparation of S expression plasmid |
| o319_S500A_Rv | GATGGCTGTCCCCGGCGGcAGAGTAGAACCCGTAATTCT TG | Preparation of S expression plasmid |
| o319_Y498A_S500A_Fw | CAAGAATTACGGGTTCgcCTCTgCCGCCGGGGACAGCCA TC | Preparation of S expression plasmid |
| o319_Y498A_S500A_Rv | GATGGCTGTCCCCGGCGGcAGAGgcGAACCCGTAATTCT TG | Preparation of S expression plasmid |
| K27A_F | GAGGACGAGGCCAAGGCCTTTTTGAACGAC | Preparation of ACE2 expression plasmid |
| K27A_R | GTCGTTCAAAAAGGCCTTGGCCTCGTCCTC | Preparation of ACE2 expression plasmid |
| D31A_F | CAAGAAATTTTTGAACGCCTTTAACTCCGAAGCTG | Preparation of ACE2 expression plasmid |
| D31A_R | CAGCTTCGGAGTTAAAGGCGTTCAAAAATTTCTTG | Preparation of ACE2 expression plasmid |
| cornutus_N38D_Fw | AACTCCGAAGCTGAAGACCTGACTTATCAAAG | Preparation of ACE2 expression plasmid |
| cornutus_N38D_Rv | CTTTGATAAGTCAGGTCTTCAGCTTCGGAGTT | Preparation of ACE2 expression plasmid |
| T40A_F | CGAAGCTGAAAACCTGGCCTATCAAAGTTCACTTG | Preparation of ACE2 expression plasmid |
| T40A_R | CAAGTGAACTTTGATAGGCCAGGTTTTCAGCTTCG | Preparation of ACE2 expression plasmid |
| Y41A_F | CTGAAAACCTGACTGCCCAAAGTTCACTTGC | Preparation of ACE2 expression plasmid |
| Y41A_R | GCAAGTGAACTTTGGGCAGTCAGGTTTTCAG | Preparation of ACE2 expression plasmid |
| Q42A_F | GAAAACCTGACTTATGCCAGTTCACTTGCTTC | Preparation of ACE2 expression plasmid |

| | | |
|---|---|---|
| Q42A_R | GAAGCAAGTGAACTGGCATAAGTCAGGTTTTC | Preparation of ACE2 expression plasmid |
| E75A_F | GTCTGCCTTTTATGAAGCCCAGTCCAAGATTGC | Preparation of ACE2 expression plasmid |
| E75A_R | GCAATCTTGGACTGGGCTTCATAAAAGGCAGAC | Preparation of ACE2 expression plasmid |
| K353A_F | GCCTGGGACCTGGGGGCCGGTGACTTCAGGATC | Preparation of ACE2 expression plasmid |
| K353A_R | GATCCTGAAGTCACCGGCCCCCAGGTCCCAGGC | Preparation of ACE2 expression plasmid |
| K27_D31A_F | TGAGGACGAGGCCAAGGCCTTTTTGAACGCCTTTAACTC CGAAGCTG | Preparation of ACE2 expression plasmid |
| K27_D31A_R | CAGCTTCGGAGTTAAAGGCGTTCAAAAAGGCCTTGGCCT CGTCCTCA | Preparation of ACE2 expression plasmid |
| T40_Y41_Q42A_F | CGAAGCTGAAAACCTGGCCGCCGCCAGTTCACTTGCTT CTTGG | Preparation of ACE2 expression plasmid |
| T40_Y41_Q42A_R | CCAAGAAGCAAGTGAACTGGCGGCGGCCAGGTTTTCAG CTTCG | Preparation of ACE2 expression plasmid |
| GD-1-2019-inf-Fw | ctatagggcgaattgGGTACCATGCTGTTCTTCTTC | Preparation of ACE2 expression plasmid |
| GD-1-2019-inf-Rv | agctccaccgcggtgGCGGCCGCTCAAGTATAGTGCAGTTT | Preparation of ACE2 expression plasmid |
| GD2019_N439K_Fw | GATTGCCTGGAACAGCAAgAACCTGGACTCTAAG | Preparation of ACE2 expression plasmid |
| GD2019_N439K_Rv | CTTAGAGTCCAGGTTcTTGCTGTTCCAGGCAATC | Preparation of ACE2 expression plasmid |
| GD2019_T478K_Fw | CTACCAGGCTGGCAGCAagCCCTGCAATGGCGTT | Preparation of ACE2 expression plasmid |
| GD2019_T478K_Rv | AACGCCATTGCAGGGctTGCTGCCAGCCTGGTAG | Preparation of ACE2 expression plasmid |
| GD2019_V483Q_Fw | CACCCCCTGCAATGGCcagGAAGGATTCAACTGC | Preparation of ACE2 expression plasmid |
| GD2019_V483Q_Rv | GCAGTTGAATCCTTCctgGCCATTGCAGGGGGTG | Preparation of ACE2 expression plasmid |
| GD2019_E484T_Fw | CCCTGCAATGGCGTTaccGGATTCAACTGCTAC | Preparation of ACE2 expression plasmid |
| GD2019_E484T_Rv | GTAGCAGTTGAATCCggtAACGCCATTGCAGGG | Preparation of ACE2 expression plasmid |
| GD2019_Q493Y_Fw | CTGCTACTTTCCTCTTtacTCATACGGCTTCCATC | Preparation of ACE2 expression plasmid |
| GD2019_Q493Y_Rv | GATGGAAGCCGTATGAgtaAAGAGGAAAGTAGCAG | Preparation of ACE2 expression plasmid |
| GD2019_H498Y_Fw | CAGTCATACGGCTTCtacCCTACCAACGGGGTG | Preparation of ACE2 expression plasmid |
| GD2019_H498Y_Rv | CACCCCGTTGGTAGGgtaGAAGCCGTATGACTG | Preparation of ACE2 expression plasmid |
| GD2019_N501D_Fw | GGCTTCCATCCTACCgACGGGGTGGGCTATCAG | Preparation of ACE2 expression plasmid |
| GD2019_N501D_Rv | CTGATAGCCCACCCCGTcGGTAGGATGGAAGCC | Preparation of ACE2 expression plasmid |
| GD2019_478-484_Fw | CTACCAGGCTGGCAGCAagCCCTGCAATGGCcagaccGG ATTCAACTGCTAC | Preparation of ACE2 expression plasmid |
| GD2019_478-484_Rv | GTAGCAGTTGAATCCggtctgGCCATTGCAGGGctTGCTGC CAGCCTGGTAG | Preparation of ACE2 expression plasmid |
| GD2019_493-501_Fw | CTGCTACTTTCCTCTTtacTCATACGGCTTCtacCCTACCgA CGGGGTGGGCTATCAG | Preparation of ACE2 expression plasmid |
| GD2019_493-501_Rv | CTGATAGCCCACCCCGTcGGTAGGgtaGAAGCCGTATGA gtaAAGAGGAAAGTAGCAG | Preparation of ACE2 expression plasmid |
| RBD_1_Fw | CAGACTAGTAACTTTAGAGTCCAGCCTACAG | Preparation of S expression plasmid |
| RBD_1_Rv | CTGTAGGCTGGACTCTAAAGTTACTAGTCTG | Preparation of S expression plasmid |
| RBD_2_Fw | CAAGTGCGTCAATTTCAATTTCAATGGGCTG | Preparation of S expression plasmid |
| RBD_2_Rv | CAGCCCATTGAAATTGAAATTGACGCACTTG | Preparation of S expression plasmid |
| MRCA2_K439N_Fw | GTTATTGCTTGGAACAGCAAcAACCTGGACGC | Preparation of S expression plasmid |
| MRCA2_K439N_Rv | GCGTCCAGGTTgTTGCTGTTCCAAGCAATAAC | Preparation of S expression plasmid |
| MRCA2_Y493Q_Fw | GCTACTACCCGCTCcAgTCCTACGGCTTTTATC | Preparation of S expression plasmid |
| MRCA2_Y493Q_Rv | GATAAAAGCCGTAGGAcTgGAGCGGGTAGTAGC | Preparation of S expression plasmid |
| MRCA2_D501N_Fw | CTTTTATCCCACTaACGGCGTTGGGTATCAG | Preparation of S expression plasmid |
| MRCA2_D501N_Rv | CTGATACCCAACGCCGTtAGTGGGATAAAAG | Preparation of S expression plasmid |
| MRCA2_K478T_Fw | CCAGGCCGGGTCCAccCCTTGCAATGGACAAAC | Preparation of S expression plasmid |
| MRCA2_K478T_Rv | GTTTGTCCATTGCAAGGggTGGACCCGGCCTGG | Preparation of S expression plasmid |
| MRCA2_Q483V_Fw | CAAACCTTGCAATGGAgtgACTGGACTAAATTGC | Preparation of S expression plasmid |
| MRCA2_Q483V_Rv | GCAATTTAGTCCAGTcacTCCATTGCAAGGTTTG | Preparation of S expression plasmid |

| | | |
|---|---|---|
| MRCA2_T484E_Fw | CCTTGCAATGGACAAgagGGACTAAATTGCTAC | Preparation of S expression plasmid |
| MRCA2_T484E_Rv | GTAGCAATTTAGTCCctcTTGTCCATTGCAAGG | Preparation of S expression plasmid |
| MRCA2_Y498H_Fw | CTACTCCTACGGCTTTcATCCCACTGACGGCGTTG | Preparation of S expression plasmid |
| MRCA2_Y498H_Rv | CAACGCCGTCAGTGGGATgAAAGCCGTAGGAGTAG | Preparation of S expression plasmid |
| MRCA2_478-484_Fw | CCAGGCCGGGTCCAccCCTTGCAATGGAgtggagGGACTAAATTGCTAC | Preparation of S expression plasmid |
| MRCA2_478-484_Rv | GTAGCAATTTAGTCCctccacTCCATTGCAAGGggTGGACCCGGCCTGG | Preparation of S expression plasmid |
| MRCA2_493-501_Fw | GCTACTACCCGCTCcAgTCCTACGGCTTTcATCCCACTaACGGCGTTGGGTATCAG | Preparation of S expression plasmid |
| MRCA2_493-501_Rv | CTGATACCCAACGCCGTtAGTGGGATgAAAGCCGTAGGAcTgGAGCGGGTAGTAGC | Preparation of S expression plasmid |
| MRCA1_N439K_Fw | GTTATTGCTTGGAACAGCAAgAACCTGGACTC | Preparation of S expression plasmid |
| MRCA1_N439K_Rv | GAGTCCAGGTTcTTGCTGTTCCAAGCAATAAC | Preparation of S expression plasmid |
| MRCA1_T478K_Fw | CCAGGCCGGGTCCAagCCTTGCAATGGAGTTG | Preparation of ACE2 expression plasmid |
| MRCA1_T478K_Rv | CAACTCCATTGCAAGGctTGGACCCGGCCTGG | Preparation of ACE2 expression plasmid |
| MRCA1_V483Q_Fw | CACTCCTTGCAATGGAcagGAGGGACTAAATTGC | Preparation of ACE2 expression plasmid |
| MRCA1_V483Q_Rv | GCAATTTAGTCCCTCctgTCCATTGCAAGGAGTG | Preparation of ACE2 expression plasmid |
| MRCA1_E484T_Fw | CCTTGCAATGGAGTTaccGGACTAAATTGCTAC | Preparation of ACE2 expression plasmid |
| MRCA1_E484T_Rv | GTAGCAATTTAGTCCggtAACTCCATTGCAAGG | Preparation of ACE2 expression plasmid |
| MRCA1_Q493Y_Fw | GCTACTACCCGCTCtAcTCCTACGGCTTTCAC | Preparation of ACE2 expression plasmid |
| MRCA1_Q493Y_Rv | GTGAAAGCCGTAGGAgTaGAGCGGGTAGTAGC | Preparation of ACE2 expression plasmid |
| MRCA1_H498Y_Fw | CAGTCCTACGGCTTTtACCCCACTAATGGCGTTG | Preparation of ACE2 expression plasmid |
| MRCA1_H498Y_Rv | CAACGCCATTAGTGGGGTaAAAGCCGTAGGACTG | Preparation of ACE2 expression plasmid |
| MRCA1_N501D_Fw | GGCTTTCACCCCACTgAcGGCGTTGGGTATCAG | Preparation of S expression plasmid |
| MRCA1_N501D_Rv | CTGATACCCAACGCCgTcAGTGGGGTGAAAGCC | Preparation of S expression plasmid |
| MRCA1_478-484_Fw | CCAGGCCGGGTCCAagCCTTGCAATGGAcagaccGGACTAAATTGCTAC | Preparation of S expression plasmid |
| MRCA1_478-484_Rv | GTAGCAATTTAGTCCggtctgTCCATTGCAAGGctTGGACCCGGCCTGG | Preparation of S expression plasmid |
| MRCA1_493-501_Fw | GCTACTACCCGCTCtAcTCCTACGGCTTTtACCCCACTgAcGGCGTTGGGTATCAG | Preparation of S expression plasmid |
| MRCA1_493-501_Rv | CTGATACCCAACGCCgTcAGTGGGGTaAAAGCCGTAGGAgTaGAGCGGGTAGTAGC | Preparation of S expression plasmid |

**Table S4. Cryo-EM data collection, refinement and validation statistics, related to Figure 3**

| | #1 Cryo-EM structure of Rc-o319 RBD/*R. cornutus* ACE2 complex (EMDB-) (PDB ) |
|---|---|
| **Data collection and processing** | |
| Magnification | Titan Krios (Thermo Fisher Scientific) |
| Voltage (kV) | 300 |
| Electron exposure (e–/Å$^2$) | 49.9 |
| Defocus range (μm) | −0.8 to −1.6 |
| Pixel size (Å) | 0.83 |
| Symmetry imposed | C1 |
| Initial particle images (no.) | 7,746,677 |
| Final particle images (no.) | 1,330,941 |
| Map resolution (Å) | 2.79 |
| FSC threshold | 0.143 |
| Map resolution range (Å) | |
| **Refinement** | |
| Initial model used (PDB code) | 8hxk |
| Model resolution (Å) | 2.8(masked)/3.0(unmasked) |
| FSC threshold | 0.5 |
| Model resolution range (Å) | |
| Map sharpening *B* factor (Å$^2$) | |
| Model composition | |
| Non-hydrogen atoms | 7060 |
| Protein residues | 866 |
| Ligands | - |
| *B* factors (Å$^2$) | |
| Protein | 11.59/187.39/62.38 |
| Ligand | - |
| R.m.s. deviations | |
| Bond lengths (Å) | 0.003 |
| Bond angles (°) | 0.544 |
| Validation | |
| MolProbity score | 1.70 |
| Clashscore | 8.05 |
| Poor rotamers (%) | 0 |
| Ramachandran plot | |
| Favored (%) | 98.84 |
| Allowed (%) | 1.16 |
| Disallowed (%) | 0 |
